# Glutamatergic and GABAergic neurons mediate distinct neurodevelopmental phenotypes of *STXBP1* encephalopathy

**DOI:** 10.1101/2021.07.13.452234

**Authors:** Joo Hyun Kim, Wu Chen, Eugene S. Chao, Hongmei Chen, Mingshan Xue

## Abstract

Heterozygous pathogenic variants in syntaxin-binding protein 1 (STXBP1, also known as MUNC18-1) cause *STXBP1* encephalopathy and are among the most frequent causes of developmental and epileptic encephalopathies and intellectual disabilities. STXBP1 is an essential protein for presynaptic neurotransmitter release, and its haploinsufficiency impairs glutamatergic and GABAergic neurotransmission. However, the mechanism underlying the broad spectrum of neurological phenotypes is poorly understood. Here we show that glutamatergic and GABAergic neurons mediate distinct disease features with few overlaps. Glutamatergic and GABAergic neurons-specific *Stxbp1* haploinsufficient mice exhibit different subsets of the cognitive and seizure phenotypes observed in the constitutive *Stxbp1* haploinsufficient mice. Developmental delay and most of the motor and psychiatric phenotypes are only recapitulated by GABAergic *Stxbp1* haploinsufficiency. Thus, the contrasting roles of excitatory and inhibitory signaling in *STXBP1* encephalopathy identify GABAergic dysfunction as a main disease mechanism and reveal the possibility to selectively modulate disease phenotypes by targeting specific neurotransmitter systems.

## Introduction

Synaptic dysfunction is a hallmark of many neurological disorders (Brose et al., 2010; Lepeta et al., 2016). Pathogenic variants in synaptic proteins may collectively account for more neurological disorders than any other functional group of proteins (Grant, 2019). More recently, an increasing number of mutations in presynaptic proteins are being discovered in neurodevelopmental disorders. These proteins are involved in different steps of the synaptic vesicle cycle–tethering, docking, priming, fusion, and recycling (Verhage and Sørensen, 2020; Bonnycastle et al., 2021; John et al., 2021; Melland et al., 2021). The clinical features of these synaptic vesicle cycle disorders are diverse, but the most prevalent features include intellectual disability, epilepsy, movement disorders, cerebral visual impairment, and psychiatric symptoms (Verhage and Sørensen, 2020; John et al., 2021). Although the molecular and cellular functions of many of these presynaptic proteins are well understood, the disease pathogeneses remain elusive. Thus, in-depth neurological studies in disease models with genetic and phenotypic accuracy are necessary to elucidate the neural mechanisms underlying neurodevelopmental deficits and develop therapeutic strategies.

Among this growing list of synaptic vesicle cycle disorders, *STXBP1* encephalopathy is the most frequent and caused by *de novo* heterozygous *STXBP1* mutations (Verhage and Sørensen, 2020; John et al., 2021). In fact, pathogenic variants in *STXBP1* are among the top 5 causes of pediatric epilepsy (Symonds and McTague, 2020) and top 10 causes of neurodevelopmental disorders (Deciphering Developmental Disorders Study, 2015; Kaplanis et al., 2020). All *STXBP1* encephalopathy patients have intellectual disability, and more than 80–90% of patients have epilepsy and motor dysfunctions (Stamberger et al., 2016; Abramov et al., 2020). Other less common clinical features include developmental delay, autistic traits, hyperactivity, anxiety, and aggressive behaviors (Stamberger et al., 2016; Suri et al., 2017). Since about 60% of the reported mutations are truncating variants (Stamberger et al., 2016; Abramov et al., 2020), haploinsufficiency is considered as the major disease mechanism, but a dominant-negative mechanism has also been proposed for a subset of missense variants (Chai et al., 2016; Guiberson et al., 2018). Several missense variants were modeled in *C. elegans*, and the mutant worms show locomotion defects and increased convulsions in response to pentylenetetrazol (Guiberson et al., 2018; Zhu et al., 2020). Haploinsufficiency was also modeled in fish and mice. Removing *stxbp1b*, one of the two *STXBP1* homologs in zebrafish, causes spontaneous electrographic seizures (Grone et al., 2016). The first three *Stxbp1* heterozygous knockout mouse models recapitulated only a subset of neurological phenotypes seen in patients, possibly due to a modest 25–50% reduction of Stxbp1 protein levels in these models (Hager et al., 2014; Miyamoto et al., 2017; Kovačević et al., 2018; Orock et al., 2018). We recently generated two new mouse *Stxbp1* null alleles, and the heterozygous mice (*Stxbp1^tm1a/+^* and *Stxbp1^tm1d/+^*) show 50% reduction in Stxbp1 protein levels in most brain regions. The two new models recapitulated nearly all features of *STXBP1* encephalopathy, as they show cognitive impairments, epileptic seizures, motor dysfunction, developmental delay, anxiety-like behaviors, hyperactivity, and aggression (Chen et al., 2020).

Mechanistically, how *STXBP1* heterozygous mutations lead to neurodevelopmental deficits remains elusive, even though it is well established that this protein is required for synaptic vesicle exocytosis in all neurons (Harrison et al., 1994; Verhage et al., 2000; Weimer et al., 2003; Grone et al., 2016), and its heterozygous null mutations impair synaptic transmission at both glutamatergic and GABAergic synapses (Toonen et al., 2006; Patzke et al., 2015; Orock et al., 2018; Miyamoto et al., 2019; Chen et al., 2020). As a first step to tackle this question, it is necessary to understand the roles of specific neurons, particularly glutamatergic and GABAergic neurons, in *STXBP1* encephalopathy pathogenesis. Furthermore, identifying the critical neuronal types can potentially provide therapeutic targets of disease. One study observed reduced survival and epileptiform discharges in GABAergic neurons-specific *Stxbp1* heterozygous knockout mice (Kovačević et al., 2018). However, other studies reported normal survival in a different line of GABAergic neurons-specific *Stxbp1* heterozygous knockout mice (Miyamoto et al., 2017) and instead observed spike-wave discharges in dorsal telencephalic glutamatergic neuron-specific heterozygous knockout mice (Miyamoto et al., 2019). Compared to the broad neurological and psychiatric impairments of *STXBP1* encephalopathy and *Stxbp1* haploinsufficient mice, the limited phenotypes of these cell type-specific *Stxbp1* heterozygous deletions seem to suggest that other neuronal types may be critical to the cognitive, motor, and psychiatric impairments. However, the efficacy and specificity of these *Stxbp1* conditional deletions were not confirmed. Thus, the significance of glutamatergic and GABAergic signaling to the disease pathogenesis is still unclear.

To address this question in a systematic manner, we generated and validated new glutamatergic and GABAergic neurons-specific *Stxbp1* haploinsufficient mice and determined their phenotypes in the three core disease domains–cognitive impairment, epilepsy, motor dysfunction–as well as psychiatric functions and general health. Our comparative study reveals that together, glutamatergic and GABAergic *Stxbp1* haploinsufficiencient mice recapitulate the vast majority of the phenotypes observed in the constitutive haploinsufficient mice. However, GABAergic neurons mediate most of the neurodevelopmental phenotypes, whereas glutamatergic neurons are critical to a smaller, but different subset of phenotypes. These results support the notion that GABAergic synaptic dysfunction is a key mechanism shared among many neurodevelopmental disorders.

## Results

### Generation of glutamatergic or GABAergic neurons-specific *Stxbp1* haploinsufficient mice

To conditionally delete *Stxbp1* in mice, we crossed a previously generated *Stxbp1^tm1a/+^* mouse (Chen et al., 2020) with a Flp recombinase germline deleter mouse (see Materials and methods) to create a new *Stxbp1* flox allele (*tm1c*), in which exon 7 is flanked by two *loxP* sites (***Figure 1A***). Heterozygous (*Stxbp1^f/+^*) and homozygous (*Stxbp1^f/f^*) flox mice are viable and fertile. Western blots with antibodies recognizing either the N- or C-terminus of Stxbp1 showed that *Stxbp1^f/+^* and *Stxbp1^f/f^* mice had similar Stxbp1 protein levels to their wild type (WT) littermates at postnatal day 0 and 3 months of age (***Figure 1B*** and ***Figure 1-supplement 1***), indicating that the presence of *FRT* and *loxP* sites does not affect Stxbp1 expression. Deletion of exon 7 by Cre recombinase will lead to an early stop codon in exon 8 (***Figure 1A***), resulting in the same *Stxbp1 tm1d* null allele described previously (Chen et al., 2020).

**Figure 1.**
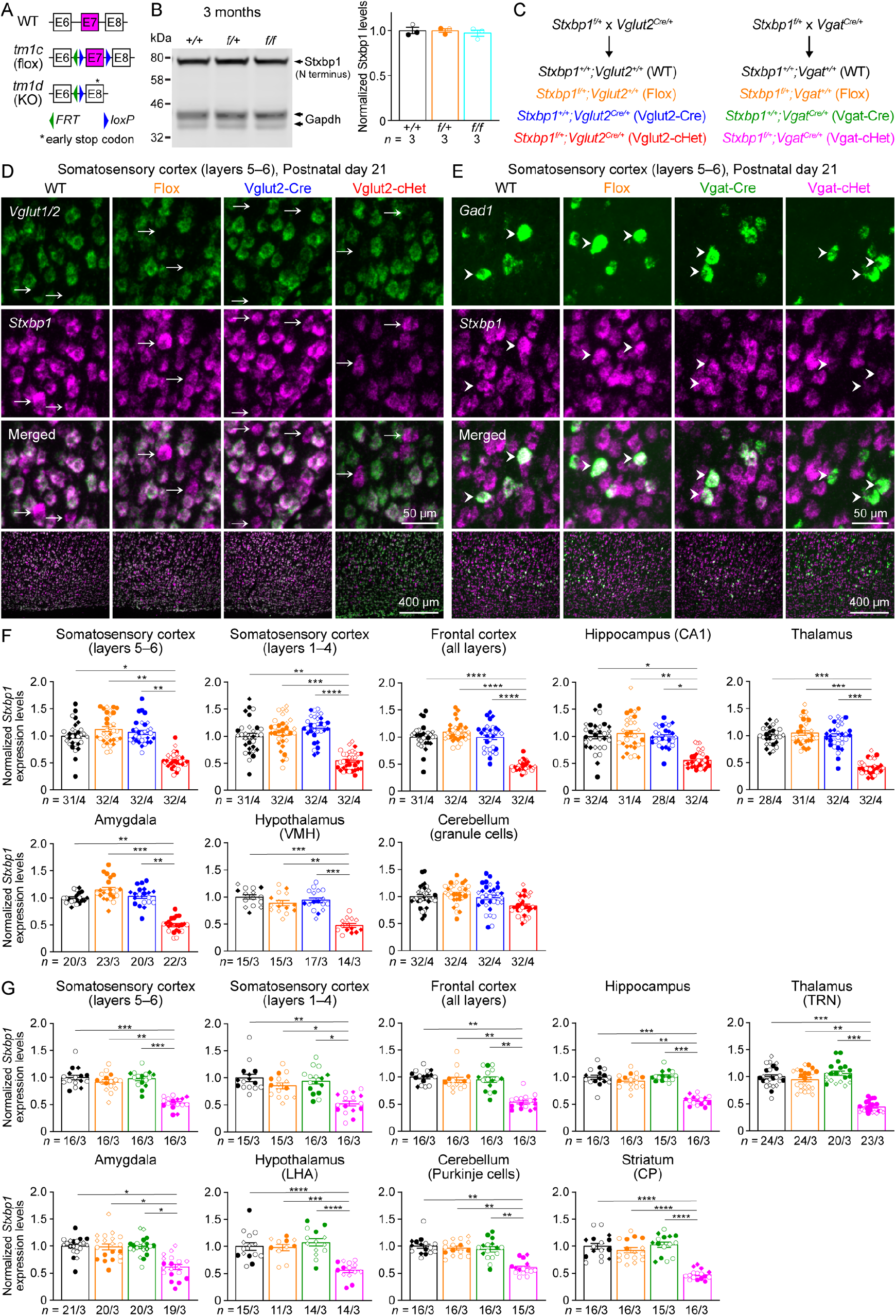
Vglut2-cHet and Vgat-cHet mice show specific reduction of *Stxbp1* levels in glutamatergic and GABAergic neurons, respectively, across different brain regions. (**A**) Genomic structures of *Stxbp1* WT, *tm1c* (flox), and *tm1d* (KO) alleles. In the flox allele, exon 7 is flanked by two *loxP* sites. In the KO allele, exon 7 is deleted, resulting in a premature stop codon in exon 8. E, exon; *FRT*, Flp recombination site; *loxP*, Cre recombination site. (**B**) Left, a representative Western blot of proteins from the cortices of 3-month-old WT, *Stxbp1^f/+^*, and *Stxbp1^f/f^* mice. Stxbp1 was detected by an antibody recognizing its N terminus. Gapdh, a housekeeping protein as loading control. Right, summary data of normalized Stxbp1 protein levels at 3 months of age. Stxbp1 levels were first normalized by the Gapdh levels and then by the average Stxbp1 levels of all WT mice from the same blot. Each filled (male) or open (female) circle represents one mouse. (**C**) *Stxbp1^f/+^* mice were crossed to *Vglut2^Cre/+^* or *Vgat^Cre/+^* mice to generate different genotypes of mice for experiments. The color scheme is maintained across all figures. (**D,E**) Representative fluorescent images from brain sections labeled by ISH probes against *Stxbp1* and *Vglut1/2* (D) or *Gad1* (E). The bottom row shows the layers 5–6 of the somatosensory cortices, and the top three rows show the individual cells from this region. Arrows (D) indicate *Vglut1/2*-negative cells, and arrow heads (E) indicates *Gad1*-positive cells. (**F**) Summary data of normalized *Stxbp1* mRNA levels in *Vglut1/2*-positive cells from different brain regions. *Stxbp1* levels were normalized by the average *Stxbp1* levels of WT brain sections that were simultaneously stained and imaged. The *Stxbp1* levels of Vglut2-cHet mice were reduced in most brain regions except cerebellar granule cells. Different shapes of symbols represent different mice (4 mice per genotype, filled circle and diamond for 2 males and open circle and diamond for 2 females), and each symbol represents one brain section. VMH, ventromedial hypothalamic nucleus. (**G**) Similar to (F), but for *Gad1*-positive cells and 3 mice per genotype. The *Stxbp1* levels of Vgat-cHet mice were reduced in all brain regions. TRN, thalamic reticular nucleus; LHA, lateral hypothalamic area; CP, caudoputamen. Bar graphs are mean ± s.e.m. * *P* < 0.05, ** *P* < 0.01, *** *P* < 0.001, **** *P* < 0.0001.

To create *Stxbp1* haploinsufficiency selectively in glutamatergic neurons, we used a pan-glutamatergic neuron-specific Cre line, *Vglut2-ires-Cre* (Vong et al., 2011). Vglut2 (vesicular glutamate transporter 2) is expressed in all glutamatergic neurons at embryonic and early postnatal stages (Boulland et al., 2004). Thus, Cre-mediated recombination occurs in all glutamatergic neurons. We crossed *Stxbp1^f/+^* mice with *Vglut2^Cre/+^* mice to obtain four genotypes: *Stxbp1^+/+^;Vglut2^+/+^* (WT), *Stxbp1^f/+^;Vglut2^+/+^* (Flox), *Stxbp1^+/+^;Vglut2^Cre/+^* (Vglut2-Cre), and *Stxbp1^f/+^;Vglut2^Cre/+^* (Vglut2-cHet). Similarly, we used a pan-GABAergic neuron-specific Cre line, *Vgat-ires-Cre* (Vong et al., 2011) to create *Stxbp1* haploinsufficiency selectively in GABAergic neurons. *Stxbp1^f/+^* mice were crossed with *Vgat^Cre/+^* mice to obtain *Stxbp1^+/+^;Vgat^+/+^* (WT), *Stxbp1^f/+^;Vgat^+/+^* (Flox), *Stxbp1^+/+^;Vgat^Cre/+^* (Vgat-Cre), and *Stxbp1^f/+^;Vgat^Cre/+^* (Vgat-cHet) (***Figure 1C***). All mice were on the C57BL/6J isogenic background.

To determine the efficiency and specificity of *Stxbp1* deletion in Vglut2-cHet and Vgat-cHet mice, we performed double fluorescent in situ hybridization (DFISH) to examine *Stxbp1* mRNA levels, as antibody staining cannot precisely identify the neuronal somas due to the presence of Stxbp1 in neurites (Ramos-Miguel et al., 2015). For Vglut2-cHet mice, we combined the probes against *Vglut1* and *Vglut2* to label all glutamatergic neurons with a single color. Compared to the three control groups (WT, Flox, and Vglut2-Cre), *Stxbp1* mRNA levels in Vglut2-cHet were reduced by 44–60% in glutamatergic neurons of most brain regions except cerebellar granular cells where the reduction was about 15–21% (***Figure 1D,F; Figure 1-supplement 2; Figure 1-supplement 3***). In contrast, *Stxbp1* mRNA levels in the *Vglut1* and *Vglut2*-negative neurons of the cortex, thalamic reticular nucleus (TRN), and striatum, which are mostly GABAergic neurons, were unaltered (***Figure 1D*; *Figure 1-supplement 2A,B*; *Figure 1-supplement 3A,B,D***). For Vgat-cHet mice, we used a probe against *Gad1* (glutamate decarboxylase 1) to label GABAergic neurons. *Stxbp1* mRNA levels in Vgat-cHet were reduced by 36–59% in GABAergic neurons of all brain regions (***Figure 1E,G; Figure 1-supplement 4*; *Figure 1-supplement 5***). Furthermore, *Stxbp1* mRNA levels in the *Gad1*-negative neurons of the cortex, thalamus, and cerebellum, which are mostly glutamatergic neurons, were unaltered (***Figure 1E*; *Figure 1-supplement 4A,B*; *Figure 1-supplement 5A,C,D***). Thus, both *Vglut2-ires-Cre* and *Vgat-ires-Cre* efficiently and specifically deleted *Stxbp1* exon 7 and caused nonsense-mediated mRNA decay. These results demonstrate that Vglut2-cHet and Vgat-cHet are indeed glutamatergic and GABAergic neurons-specific *Stxbp1* haploinsufficient mice, respectively.

Guided by the phenotypes of constitutive haploinsufficient mice *Stxbp1^tm1d/+^* (Chen et al., 2020) and symptoms of *STXBP1* encephalopathy patients, we sought to characterize the neurological functions of male and female Vglut2-cHet and Vgat-cHet mice in comparison with their sex- and age-matched control littermates to dissect the contributions of glutamatergic and GABAergic neurons to *STXBP1* encephalopathy pathogenesis. We will conclude that glutamatergic or GABAergic neurons-specific *Stxbp1* haploinsufficiency significantly alters a phenotype only if Vglut2-cHet or Vgat-cHet mice, respectively, are statistically different from at least both Flox and Cre control mice.

### GABAergic neurons-specific *Stxbp1* haploinsufficiency causes reduced survival, reduced body weight, dystonia, and impaired nesting behavior

*Stxbp1^tm1d/+^* mice show reduced survival and body weight, and develop dystonia that manifests as hindlimb stiffness or clasping (Chen et al., 2020). Thus, we monitored Vglut2-cHet, Vgat-cHet, and their respective control mice weekly for their survival, body weight, and dystonia. Vglut2-cHet mice were observed at the expected Mendelian frequency (***Figure 2A,B***), and they had a normal survival rate when monitored up to one year (***Figure 2C,D***). Vgat-cHet mice were also observed at the expected Mendelian frequency at postnatal day 14 (***Figure 2A***), but about 20% of them died during postnatal weeks 3 and 4 (***Figure 2B***), indicating a postnatal lethality phenotype. Interestingly, those Vgat-cHet mice that survived through this period had a similar survival rate as control mice (***Figure 2C,D***). The partially penetrant lethality phenotype of Vgat-cHet mice is similar that of *Stxbp1^tm1d/+^* mice (Chen et al., 2020).

**Figure 2.**
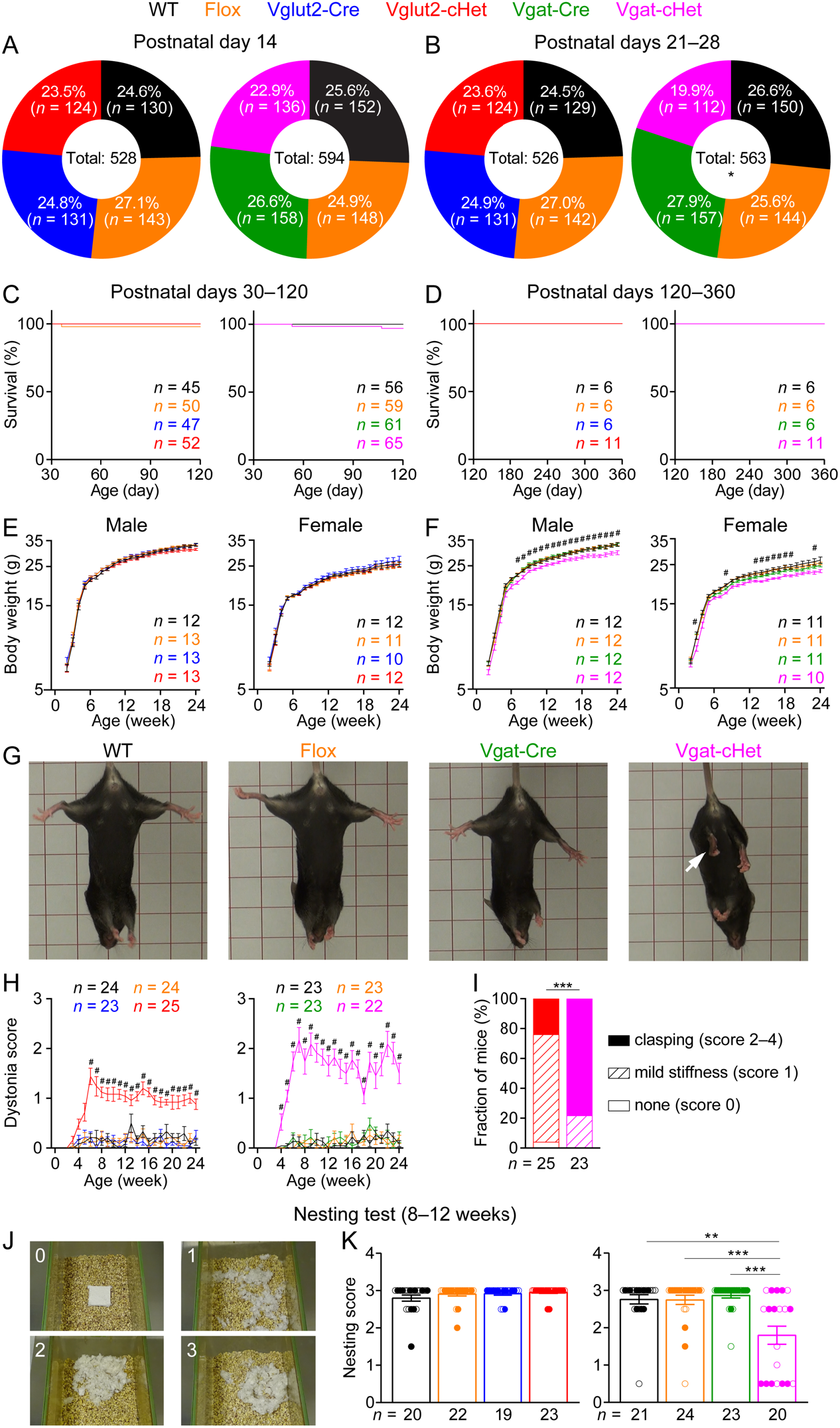
Vgat-cHet mice show reduced survival, reduced body weight, dystonia, and impaired nesting behavior. (**A,B**) Pie charts show the observed genotypes of different genotypes at postnatal day 14 (A) and postnatal days 21–28 (B). Vgat-cHet mice were significantly less than Mendelian expectations at postnatal days 21–28. (**C,D**) Both Vglut2-cHet (C) and Vgat-cHet (D) mice had normal survival rates after postnatal day 30. (**E,F**) Body weight as a function of age. The body weight of Vgat-cHet mice was less than that of control mice (F). # indicates that Vgat-cHet mice are statistically different (i.e., at least *P* < 0.05) from at least both Flox and Vgat-Cre mice. (**G**) Vgat-cHet showed dystonia and hindlimb clasping (arrows). (**H**) Dystonia scores as a function of age. # indicates that Vglut2-cHet and Vgat-cHet mice are statistically different (i.e., at least *P* < 0.05) from at least both corresponding Flox and Cre mice. (**I**) The fractions of Vglut2-cHet and Vgat-cHet mice with different severities of dystonia. (**J,K**) The quality of the nests was scored according to the criteria in (J). Vgat-cHet mice built poor quality nests within 24 hours. The numbers and ages of tested mice are indicated in the figures. Each filled (male) or open (female) circle represents one mouse. Bar graphs are mean ± s.e.m. * *P* < 0.05, ** *P* < 0.01, *** *P* < 0.001, **** *P* < 0.0001.

While the body weight of Vglut2-cHet mice was indistinguishable from that of their sex- and age-matched control littermates (***Figure 2E***), Vgat-cHet mice were smaller and weighted about 9% less than control mice (***Figure 2F***). Furthermore, both Vglut2-cHet and Vgat-cHet mice began to develop dystonia around 3–4 weeks of age and progressively exacerbated over the next month (***Figure 2G,H***). The dystonia of Vgat-cHet mice was much more severe than that of Vglut2-cHet mice. By the age of 6 months, 78% Vgat-cHet mice developed hindlimb clasping, but only 24% Vglut2-cHet mice did (***Figure 2I***).

To further assess mouse well-being, we examined the innate nest building behavior by providing a Nestlet (pressed cotton square) to each mouse in the home cage and scoring the degree of shredding and nest quality after 24 hours (***Figure 2J,K***). Vgat-cHet mice built worse nests than control mice, similar to *Stxbp1^tm1d/+^* mice (Chen et al., 2020), whereas Vglut2-cHet mice were normal (***Figure 2K***).

Altogether, these results show that *Stxbp1* haploinsufficiency in GABAergic neurons alone is sufficient to recapitulate the reduced survival, smaller body weight, dystonia, and impaired nesting behavior observed in the constitutive *Stxbp1* haploinsufficient mice.

### Increased anxiety-like behaviors and hyperactivity of GABAergic neurons-specific *Stxbp1* haploinsufficient mice

Anxiety was reported in a subset of *STXBP1* encephalopathy patients (Marchese et al., 2016; Suri et al., 2017), and several lines of constitutive *Stxbp1* heterozygous knockout mice including *Stxbp1^tm1d/+^* show increased anxiety (Hager et al., 2014; Miyamoto et al., 2017; Kovačević et al., 2018; Chen et al., 2020). Thus, we used the elevated plus maze test to assess anxiety-like behaviors. Vgat-cHet mice entered the open arms less frequently, spent less time, and traveled shorter distance in the open arms than control mice (***Figure 3A–C***), but their activities in the closed arms were largely normal (***Figure 3D***), showing the heightened anxiety of Vgat-cHet mice. In contrast, Vglut2-cHet mice did not show increased anxiety-like behaviors in this test (***Figure 3A–D***).

**Figure 3.**
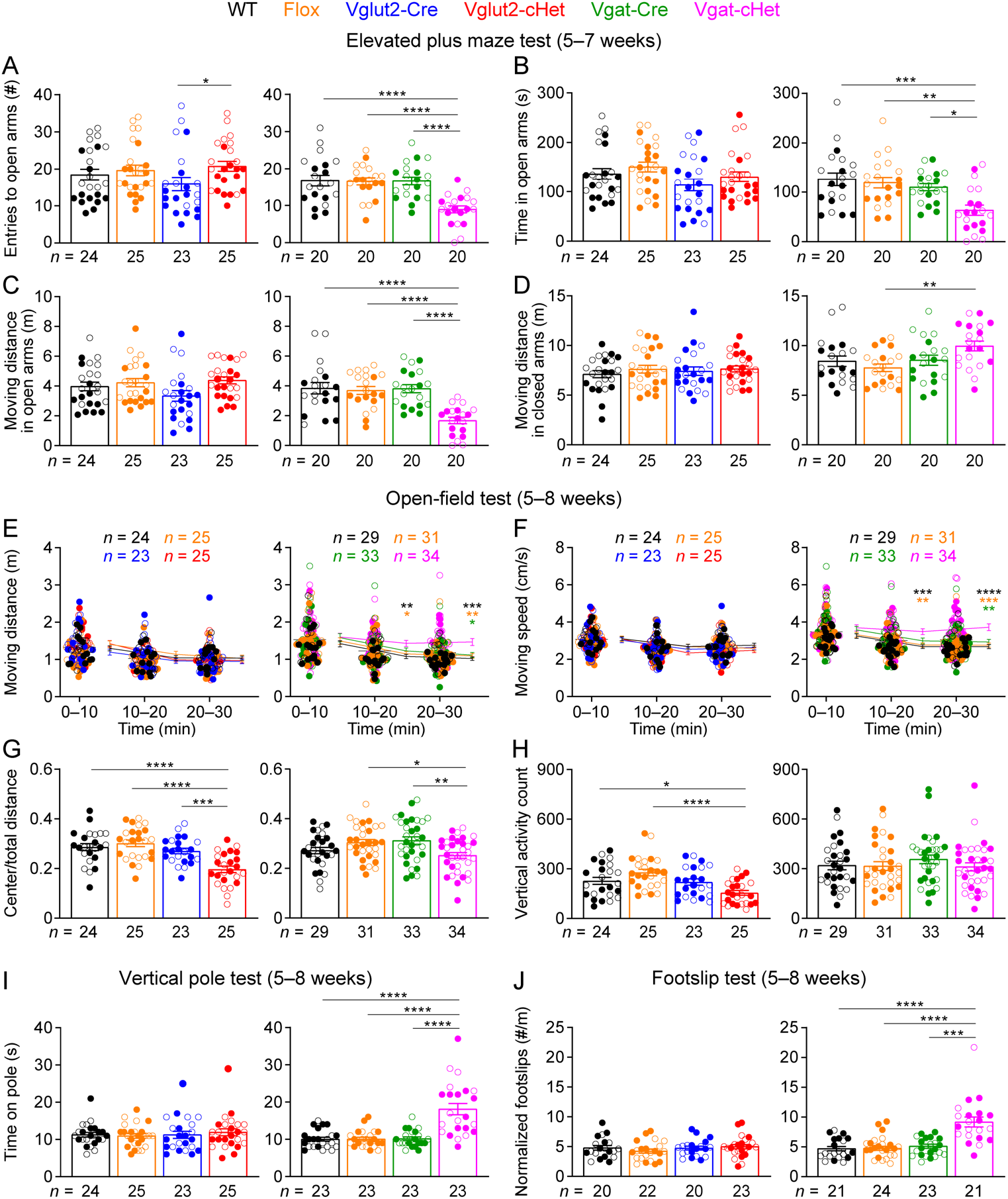
Vgat-cHet mice show elevated anxiety-like behaviors, hyperactivity, and motor dysfunctions. (**A**–**D**) In the elevated plus maze test, Vgat-cHet mice, but not Vglut2-cHet mice, entered the open arms less frequently (A), spent less time (B), and traveled shorter distance (C) in the open arms than control mice. In the closed arms, the travel distances of Vglut2-cHet mice were similar to those of control mice, and Vgat-cHet mice traveled slightly longer distances than Flox mice (D). (**E,F**) In the open-field test, Vgat-cHet mice, but not Vglut2-cHet mice, showed an increase in the moving distance (E) and speed (F). The statistical significance between Vgat-cHet and WT, Flox, or Vgat-Cre mice is indicated by black, orange, or green asterisks, respectively. (**G**) Vglut2-cHet and Vgat-cHet mice showed a decrease in the ratio of center moving distances over total moving distance. (**H**) Vglut2-cHet mice, but not Vgat-cHet mice, showed a decrease in the vertical activity. (**I,J**) Vgat-cHet mice, but not Vglut2-cHet mice, took more time to come down from a vertical pole (I) and made more foot slips per travel distance on a wire grid (J). The numbers and ages of tested mice are indicated in the figures. Each filled (male) or open (female) circle represents one mouse. Bar graphs are mean ± s.e.m. * *P* < 0.05, ** *P* < 0.01, *** *P* < 0.001, **** *P* < 0.0001.

We next used the open-field test to examine mouse locomotion, exploration, and anxiety-like behaviors, as *Stxbp1^tm1d/+^* mice show hyperactivity and increased anxiety-like behaviors in this test (Chen et al., 2020). Typically, the activities of mice decrease over time in this test, as mice become acclimated to the test arena. The locomotion of Vglut2-cHet mice was similar to that of control mice (***Figure 3E,F***). In contrast, Vgat-cHet mice showed a hyperactive phenotype, as their activities did not decrease over time and they traveled longer distances and faster than control mice in the later phase of the test (***Figure 3E,F***). Vgat-cHet mice explored the center region of the arena less than control mice (***Figure 3G***), consistent with their heightened anxiety. Interestingly, we observed that Vglut2-cHet mice also avoided the arena center region and made fewer vertical movements (***Figure 3G,H***), indicating an increase in anxiety as well.

We also performed marble burying test to evaluate innate digging behavior. The numbers of marbles buried by Vglut2-cHet or Vgat-cHet mice were not statistically different from those of the control mice, even though Vgat-cHet mice buried fewer marbles (***Figure 3-supplement 1A***). Taken the results of the elevated plus maze and open-field tests together, both glutamatergic and GABAergic neurons contribute to anxiety-like behaviors with GABAergic neurons being more critical, as elevated plus maze test is a more specific assay for anxiety-like behaviors in mice. Furthermore, *Stxbp1* haploinsufficiency in GABAergic neurons mediates the hyperactivity phenotype of constitutive *Stxbp1* haploinsufficient mice.

### Impaired motor coordination and normal sensory functions of GABAergic neurons-specific *Stxbp1* haploinsufficient mice

We next evaluated motor functions using rotarod, vertical pole, and footslip tests, as motor deficits are prevalent in *STXBP1* encephalopathy patients (Stamberger et al., 2016) and were observed in *Stxbp1^tm1d/+^* mice (Chen et al., 2020). We performed rotarod test for 4 consecutive days to evaluate motor learning and coordination by measuring the latency of mice to fall from a rotating rod. Vglut2-cHet mice performed similarly to control mice except the first trial where they fell off the rotating rod earlier (***Figure 3-supplement 2B***), whereas Vgat-cHet mice stayed longer on the rod than control mice across multiple trials (***Figure 3-supplement 2C***), probably due to their smaller body weight (***Figure 2F***) and hyperactivity (***Figure 3E,F***). This rotarod phenotype of Vgat-cHet mice is similar to that of *Stxbp1^tm1d/+^* mice (Chen et al., 2020).

The vertical pole test assesses the agility of mice by measuring the amount of time it takes for mice to descend from the top of a vertical pole. Vgat-cHet mice took 80% more time to complete this task than control mice (***Figure 3I***). When allowed to walk on a wire grid, Vgat-cHet mice had difficulty in placing their paws precisely on the wire to hold themselves and slipped 85–97% more frequently than control mice (***Figure 3J***). In contrast, Vglut2-cHet mice performed normally in both vertical pole and footslip tests (***Figure 3I,J***). Together, these results show that like constitutive *Stxbp1* haploinsufficient mice, Vgat-cHet mice do not develop ataxia, but their fine motor coordination is impaired.

Finally, both Vglut2-cHet and Vgat-cHet mice showed normal acoustic startle responses (***Figure 3-supplement 2A***), pre-pulse inhibition (***Figure 3-supplement 2B***), and thermal nociception (***Figure 3-supplement 2C***). Since *Stxbp1^tm1d/+^* mice are normal in these tests too, these results indicate that glutamatergic or GABAergic neurons-specific *Stxbp1* haploinsufficiency does not lead to additional abnormalities in sensory functions and sensorimotor gating.

### Social aggression is increased in GABAergic neurons-specific *Stxbp1* haploinsufficient mice

A subset of *STXBP1* encephalopathy patients exhibit autistic features and aggressive behaviors (Stamberger et al., 2016; Abramov et al., 2020). *Stxbp1^tm1d/+^* mice show normal social interactions in the three-chamber test, but male resident *Stxbp1^tm1d/+^* mice exhibit elevated innate aggression towards male intruder mice in the resident-intruder test (Chen et al., 2020). Thus, we evaluated Vglut2-cHet and Vgat-cHet mice in these two tests. In the three-chamber test, both Vglut2-cHet and Vgat-cHet mice preferred to interact with a sex- and age-matched partner mouse rather than an object, similar to the control mice (***Figure 4A***), showing their normal sociability. In the resident-intruder test, male resident Vgat-cHet mice started the first attack sooner, initiated more attacks, and spent more time attacking the intruders than control mice (***Figure 4B–E***), all of which indicate an elevated innate aggression. In contrast, Vglut2-cHet mice were not statistically different from control mice in any of these parameters, although there might be signs of elevated aggression based on the number of attacks and total duration of attacks (***Figure 4B–E***). These results indicate that GABAergic neurons are critically involved in the elevated innate aggression caused by *Stxbp1* haploinsufficiency.

**Figure 4.**
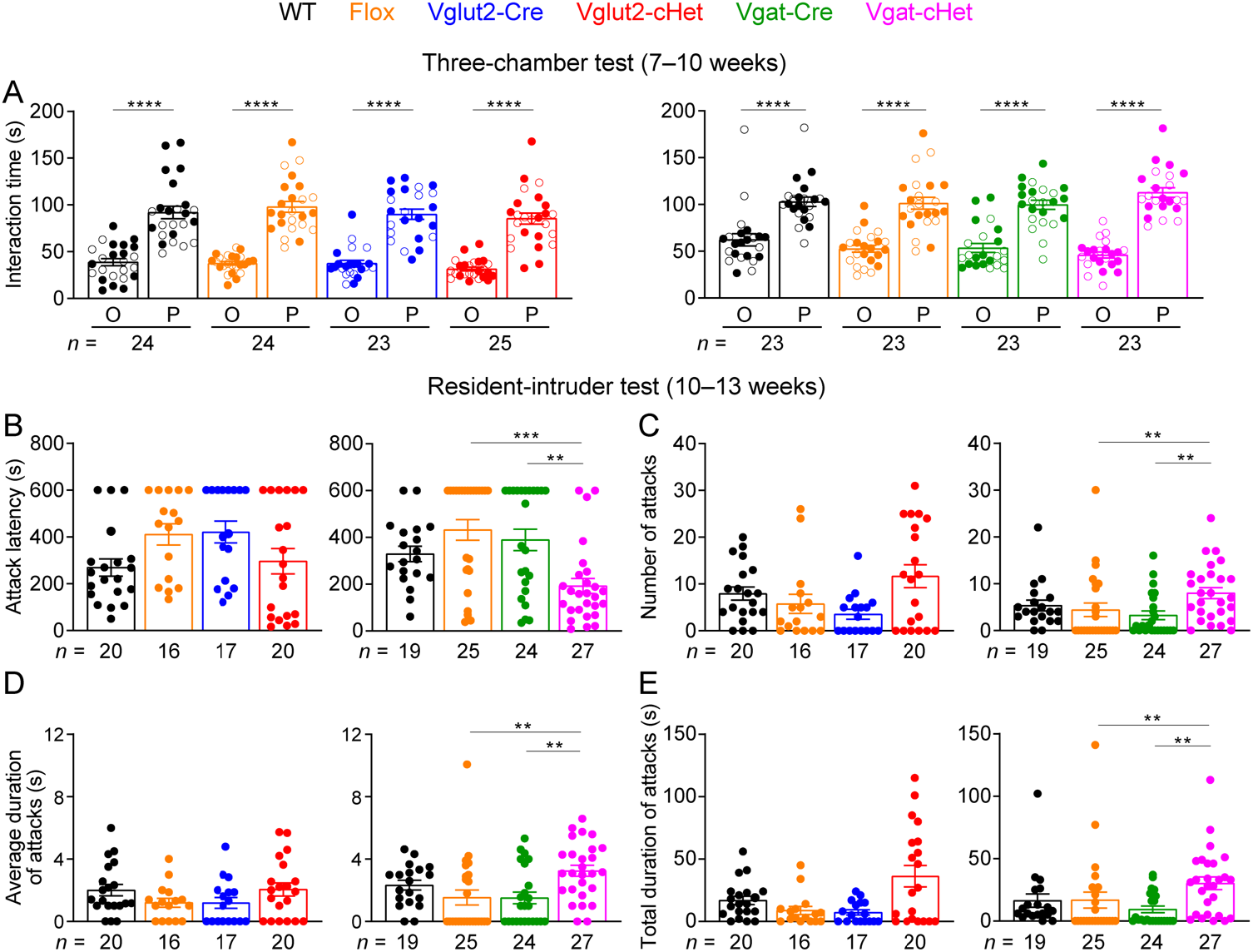
Vgat-cHet mice show normal sociability but increased aggressive behaviors. (**A**) In the three-chamber test, Vglut2-cHet, Vgat-cHet, and control mice showed a preference in interacting with the partner mouse over the object. (**B**–**E**) In the resident-intruder test, male Vgat-cHet mice, but not Vglut2-cHet mice, showed a reduction in the latency to attack the male intruder mice (B). The number (C), average duration (D), and total duration (E) of attacks were increased as compared to control mice. The numbers and ages of tested mice are indicated in the figures. Each filled (male) or open (female) circle represents one mouse. Bar graphs are mean ± s.e.m. ** *P* < 0.01, *** *P* < 0.001, **** *P* < 0.0001.

### Glutamatergic and GABAergic neurons-specific *Stxbp1* haploinsufficiencies differentially impair cognitive functions

One of the core features of *STXBP1* encephalopathy is intellectual disability (Stamberger et al., 2016; Abramov et al., 2020), which is recapitulated by the severe cognitive deficits in *Stxbp1^tm1d/+^* mice (Chen et al., 2020). Thus, we assessed the cognitive functions of Vglut2-cHet and Vgat-cHet mice. We first performed the novel object recognition test, in which WT mice prefer to explore a novel object over a familiar object, whereas *Stxbp1^tm1d/+^* mice fails to recognize the novel object (Chen et al., 2020). Surprisingly, neither Vglut2-cHet nor Vgat-cHet mice showed a deficit in this test (***Figure 5A; Figure 5-supplement 1***). This result was unexpected because novel object recognition is thought to depend on the hippocampus and cortex (Antunes and Biala, 2012; Cohen and Stackman, 2015), and *Stxbp1* haploinsufficiency impairs GABAergic synaptic transmission in the cortex (Chen et al., 2020), which is expected to alter cortical functions in Vgat-cHet mice. Vgat-cHet mice so far have recapitulated most of the phenotypes of *Stxbp1^tm1d/+^* mice. Thus, the intact recognition memories in Vglut2-cHet and Vgat-cHet mice indicate that *Stxbp1* haploinsufficiency in glutamatergic or GABAergic neurons alone is not sufficient to impair novel object recognition or other neuronal types are more important for this cognitive function.

**Figure 5.**
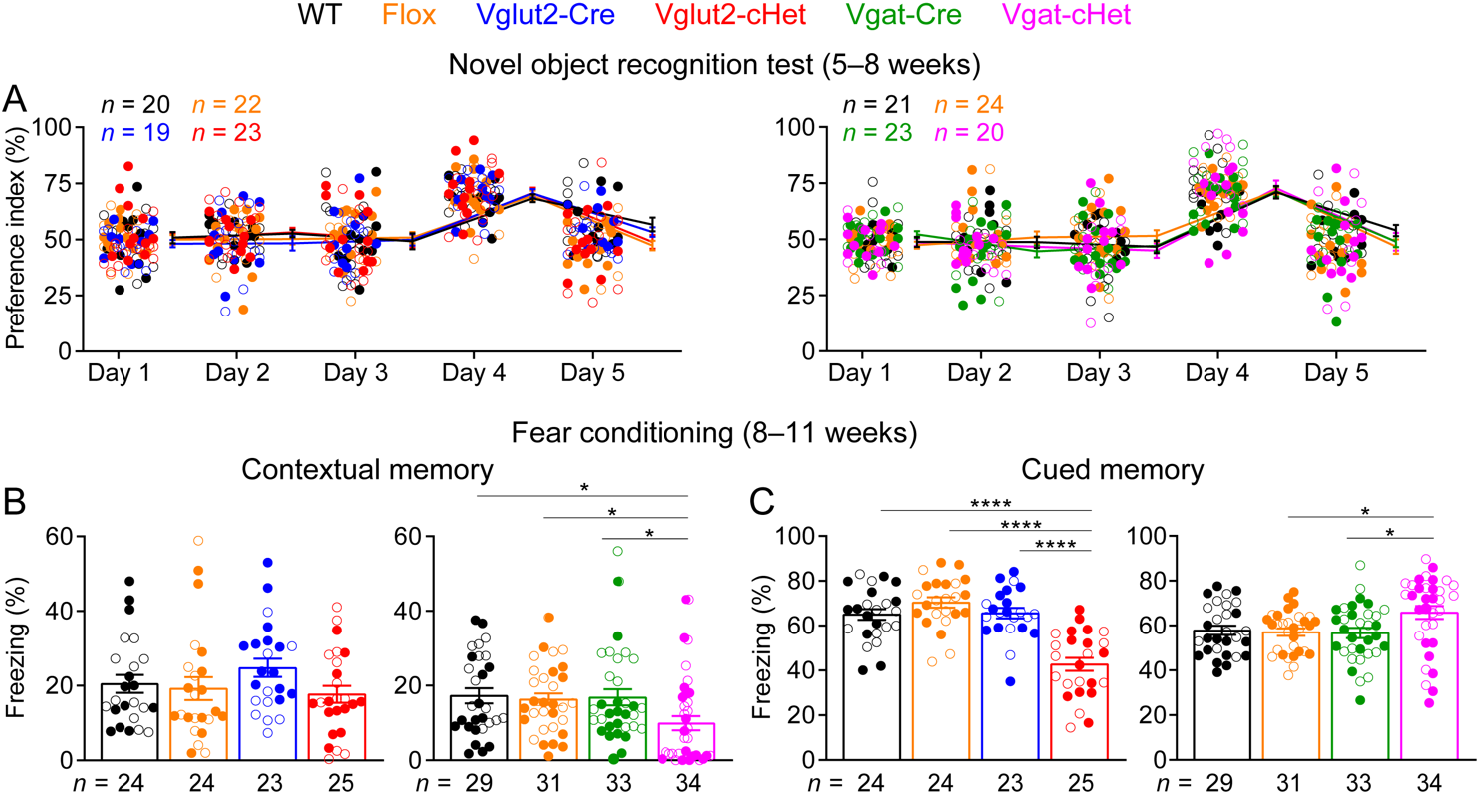
Distinct cognitive deficits of Vglut2-cHet and Vgat-cHet mice. (**A**) In the novel object recognition test with 24-hour testing intervals, the ability of a mouse to recognize the novel object was measured by the preference index (see Materials and methods). Mice were presented with the same two identical objects on days 1, 2, 3, and 5, and the familiar object and a novel object on day 4. Similar to control mice, Vglut2-cHet and Vgat-cHet mice showed a preference for the novel object. (**B,C**) In the fear conditioning test, Vgat-cHet and Vglut2-cHet mice showed a reduction in context-induced (B) and cue-induced (C) freezing behaviors 24 hours after training, respectively. Vgat-cHet also showed a modest increase in cue-induced freezing. The numbers and ages of tested mice are indicated in the figures. Each filled (male) or open (female) circle represents one mouse. Bar graphs are mean ± s.e.m. * *P* < 0.05, **** *P* < 0.0001.

To further examine cognitive functions by another test, we evaluated Vglut2-cHet and Vgat-cHet mice in the Pavlovian fear conditioning paradigm, in which *Stxbp1^tm1d/+^* mice display a strong reduction in both context- and cue-induced freezing behaviors 24 hours after conditioning (Chen et al., 2020). Interestingly, Vglut2-cHet and Vgat-cHet mice showed a selective deficit in hippocampus-independent cued fear memory and hippocampus-dependent contextual fear memory, respectively (***Figure 5B,C***). Vglut2-cHet mice were normal in contextual memory (***Figure 5B***), whereas Vgat-cHet mice even had slightly better cued memory than control mice (***Figure 5C***). The reduced freezing responses in Vglut2-cHet and Vgat-cHet mice were not due to sensory dysfunctions as their acoustic startle responses and nociception were intact (***Figure 3-supplement 3A,C***). This striking segregation of two forms of associative memory in Vglut2-cHet and Vgat-cHet mice highlights the importance of both glutamatergic and GABAergic neurons in the cognitive deficits of *STXBP1* encephalopathy.

### Distinct epileptic seizures in glutamatergic and GABAergic neurons-specific *Stxbp1* haploinsufficient mice

Epilepsy is a hallmark feature of *STXBP1* encephalopathy and patients present diverse types including epileptic spasm, focal, tonic, clonic, myoclonic, and absence seizures (Stamberger et al., 2016; Suri et al., 2017). Constitutive *Stxbp1* heterozygous knockout mice including *Stxbp1^tm1d/+^* mice have frequent spike-wave discharges (SWDs) and myoclonic seizures that manifested as involuntary muscle jerks associated with EEG discharges or sudden jumps (Kovačević et al., 2018; Miyamoto et al., 2019; Chen et al., 2020). We performed chronic video-electroencephalography (EEG) and electromyography (EMG) recordings in freely moving Vglut2-cHet, Vgat-cHet, and control mice (***Figure 6A–C***). All Vglut2-cHet mice exhibited numerous spike-wave discharges (SWDs), which is similar to *Stxbp1^tm1d/+^* mice (***Figure 6B,D; Video 1***). This result is also consistent with the report that heterozygous deletion of *Stxbp1* in dorsal telencephalic glutamatergic neurons caused frequent SWDs (Miyamoto et al., 2019). The myoclonic jerks and jumps of Vglut2-cHet mice were indistinguishable from those of control mice (***Figure 6E,G***). In contrast, Vgat-cHet mice did not show frequent SWDs (***Figure 6D***), but all had many myoclonic jerks and jumps, particularly during rapid eye movement (REM) and non-rapid eye movement (NREM) sleeps (***Figure 6C,E–G; Video 2; Video 3***). Thus, the segregation of two types of seizures in Vglut2-cHet and Vgat-cHet mice again highlights the important, but different, roles of glutamatergic and GABAergic neurons in the epileptogenesis for *STXBP1* encephalopathy.

**Figure 6.**
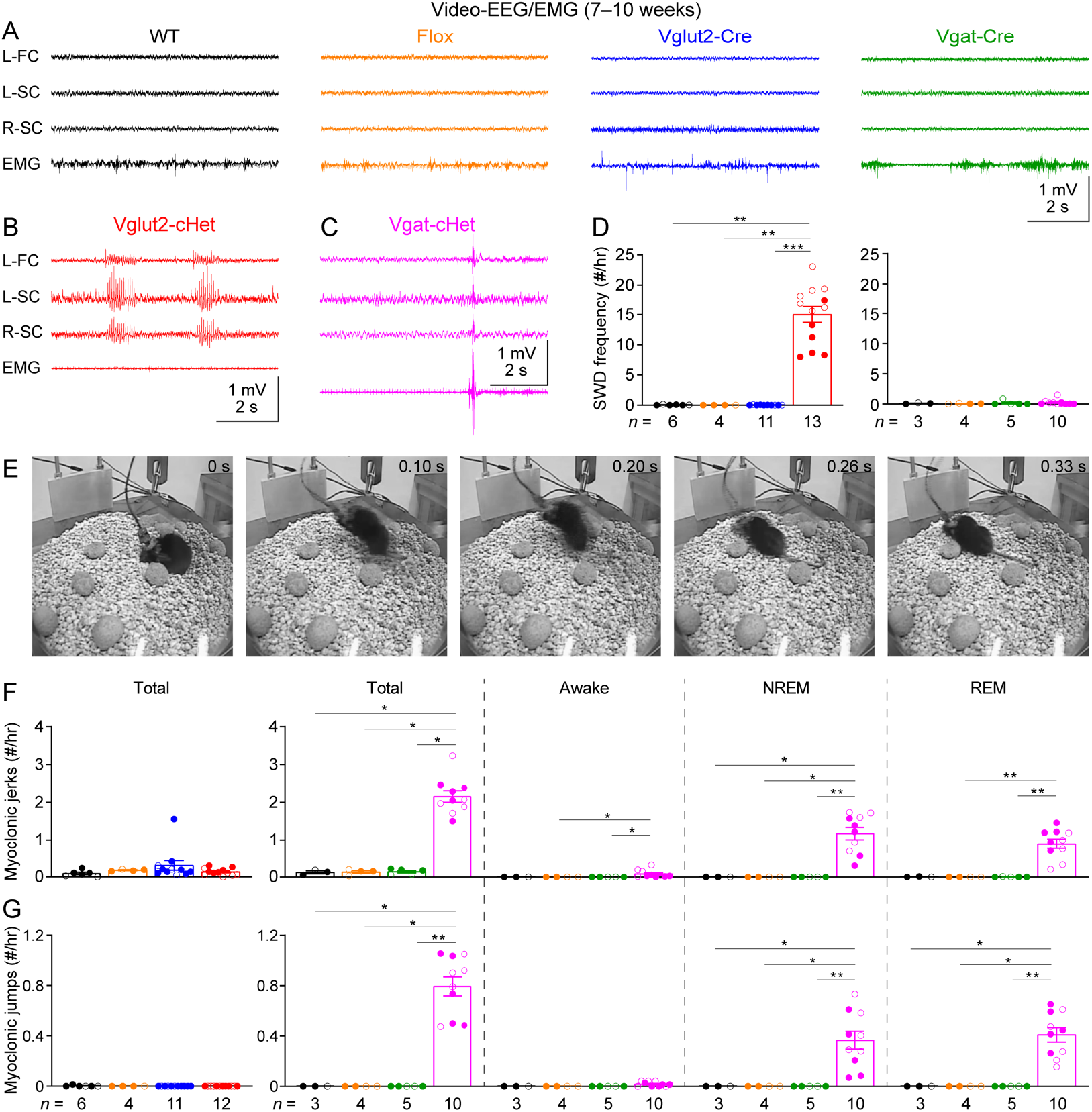
Vglut2-cHet and Vgat-cHet mice exhibit different forms of epileptic seizures. (**A**) Representative EEG traces from the left frontal cortex (L-FC), left somatosensory cortex (L-SC), right somatosensory cortex (R-SC), and EMG traces from the neck muscle of control mice. (**B,C**) Representative EEG and EMG traces showing two spike-wave discharges (SWDs, indicated by the blue arrows) of a Vglut2-cHet mouse (B) and a myoclonic jerk (indicated by the blue arrow) of a Vgat-cHet mouse (C). The Vgat-cHet mouse was in REM sleep before the jerk. (**D**) Summary data showing that the SWD frequencies of Vglut2-cHet mice were drastically increased as compared to control mice. (**E**) Video frames showing a myoclonic jump of a Vgat-cHet mouse (see ***Video 3***). The mouse was in REM sleep before the jump. (**F,G**) Summary data showing the total frequencies of myoclonic jerks (F) and jumps (G) and the frequencies in different behavioral states. The frequencies of both jerks and jumps were drastically increased in Vgat-cHet mice, particularly during NREM and REM sleep. The numbers and ages of recorded mice are indicated in the figures. Each filled (male) or open (female) circle represents one mouse. Bar graphs are mean ± s.e.m. * *P* < 0.05, ** *P* < 0.01, *** *P* < 0.001.

## Discussion

Glutamatergic and GABAergic neurons together mediate most of the neurological impairments of *STXBP1* encephalopathy. Vgat-cHet mice exhibit reduced body weight, early lethality, hindlimb clasping, motor dysfunction, impaired nest building, hyperactivity, aggression, impaired contextual fear memory, and myoclonic seizures, whereas Vglut2-cHet mice show impaired cued fear memory and SWDs. Only the impaired marble burying and novel object recognition of constitutive *Stxbp1* haploinsufficient mice are not recapitulated by either Vglut2-cHet or Vgat-cHet mice (***Figure 7***; ***Supplementary File 2***). Thus, it is likely that glutamatergic and especially GABAergic neurons are the most critical cell types to *STXBP1* encephalopathy. It would be worth determining to what extent the neurodevelopmental deficits of constitutive *Stxbp1* haploinsufficient mice can be rescued by restoring *Stxbp1* expression solely in GABAergic neurons. We previously observed two deficits of cortical inhibition in constitutive *Stxbp1* haploinsufficient mice – a reduction in the synaptic strength of parvalbumin-expressing interneurons and a reduction in the connectivity of somatostatin-expressing interneurons (Chen et al., 2020). It would be interesting to determine the specific contribution of each of these two main classes of GABAergic neurons to the disease pathogenesis. Future experiments are also necessary to determine if other neurotransmitter systems such as dopaminergic and noradrenergic neurons are important, particularly for object recognition memory.

**Figure 7.**
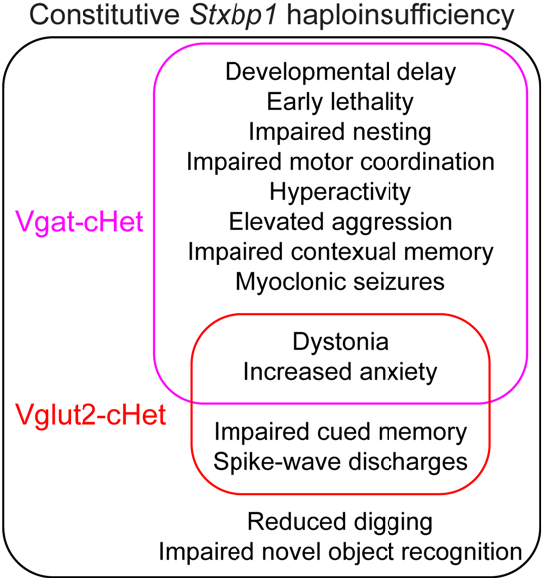
Comparison of constitutive *Stxbp1* haploinsufficient mice, Vglut2-cHet mice, and Vgat-cHet mice. Square Venn diagram showing the phenotypes of constitutive *Stxbp1* haploinsufficient mice, Vglut2-cHet mice, and Vgat-cHet mice. Except the reduced digging behavior and impaired novel object recognition, Vglut2-cHet and Vgat-cHet mice together recapitulate all other phenotypes of constitutive haploinsufficient mice. Vglut2-cHet and Vgat-cHet mice each recapitulate distinct subsets of the phenotypes of constitutive haploinsufficient mice. Only dystonia and increased anxiety are shared between Vglut2-cHet and Vgat-cHet mice. Vgat-cHet mice exhibit broader and more severe phenotypes than Vglut2-cHet mice.

GABAergic *Stxbp1* haploinsufficiency overall causes substantially broader and more severe phenotypes than glutamatergic *Stxbp1* haploinsufficiency (***Figure 7***; ***Supplementary File 2***), supporting the notion that reduced inhibition is a primary mechanism of *STXBP1* encephalopathy (Chen et al., 2020). This is in sharp contrast with previous studies that indicate a major role of impaired excitatory synaptic transmission in the disease pathogenesis (Patzke et al., 2015; Miyamoto et al., 2017; Orock et al., 2018; Miyamoto et al., 2019), in part because previous models of heterozygous deletion of *Stxbp1* in GABAergic neurons revealed few phenotypes. The first model using an *Stxbp1* exon 3-floxed allele and a bacterial artificial chromosome transgenic *Vgat-Cre* line resulted in normal survival, locomotion, fear memory, and innate aggression (Miyamoto et al., 2017). The second model using an *Stxbp1* exon 2-floxed allele and a *Gad2-ires-Cre* line showed partial early lethality, but other neurological functions were not studied (Kovačević et al., 2018). Both models had epileptiform activities that are likely the same myoclonic seizures of our model, but the lack of quantitative characterization precludes a comparison with our results in detail (Kovačević et al., 2018; Miyamoto et al., 2019). Apart from different experimental conditions or assays that may have contributed to the difference among studies, one possible difference among these models is the efficiency and specificity of *Stxbp1* deletion due to different *Cre* lines and recombination efficacies of different *Stxbp1* flox alleles, as previous studies did not validate the genetic manipulations like what we did. Since there are milder phenotypes and Stxbp1 protein reductions in the constitutive *Stxbp1* heterozygous knockout mice derived from the two previous flox alleles than our *Stxbp1* flox allele-derived *Stxbp1^tm1d/+^* mice (Miyamoto et al., 2017; Kovačević et al., 2018; Chen et al., 2020), different levels of Stxbp1 protein reduction in different *Stxbp1* flox alleles-mediated deletions may also contribute to the phenotypic differences. Furthermore, we studied Vglut2-cHet and Vgat-cHet mice in parallel and with much larger cohorts of mice than previous studies, allowing a better detection and comparison of neurodevelopmental phenotypes.

Glutamatergic and GABAergic neurons are extensively interconnected throughout the brain, and their synaptic interactions control the spatiotemporal patterns of neuronal activity and brain functions. Thus, one would expect that manipulating a gene that is important for both glutamatergic and GABAergic neurons in either neuronal type should lead to largely overlapping neurological phenotypes. For instance, deletion of Rett syndrome gene *Mecp2* (*methyl-CpG-binding protein 2*) in glutamatergic or GABAergic neurons causes many common phenotypes in mice (Gemelli et al., 2006; Chao et al., 2010; Meng et al., 2016). On the contrary, our cell type-specific deletion studies reveal distinct roles of these two neuronal types in the phenotypic spectrum of *STXBP1* encephalopathy. The only shared phenotypes between Vglut2-cHet and Vgat-cHet mice are dystonia and increased anxiety-like behaviors (***Figure 7***; ***Supplementary File 2***). Intriguingly, glutamatergic and GABAergic neurons each independently subserves one of the two forms of associative memories in Pavlovian fear conditioning and one of the two seizure types. This unexpected phenotypic segregation between excitatory and inhibitory neurons suggests that different neurological functions exhibit different susceptibilities to the presynaptic dysfunction caused by *Stxbp1* haploinsufficiency in glutamatergic or GABAergic neurons. The distinct roles of glutamatergic and GABAergic neurotransmitter systems in *STXBP1* encephalopathy present both challenges and opportunities for therapeutic interventions. Both neuronal types should be the primary targets, but their wide distribution throughout the brain makes it difficult for gene-based therapies such as adeno-associated virus-mediated gene replacement to achieve a high degree of coverage for both populations. On the other hand, the clinical symptoms vary considerably among patients, and some patients present only a subset of disease phenotypes. Thus, modulating one of these two neurotransmitter systems by small molecules such as transmitter receptor modulators may allow more precise treatment of the symptoms.

The reduction of glutamatergic and GABAergic synaptic transmission caused by *STXBP1* or *Stxbp1* heterozygous mutations are rather modest (Toonen et al., 2006; Patzke et al., 2015; Orock et al., 2018; Miyamoto et al., 2019; Chen et al., 2020), yet the neurological impairments in humans and mice are severe (Stamberger et al., 2016; Abramov et al., 2020; Chen et al., 2020), highlighting the profound impacts of subtle presynaptic dysfunctions on neuronal functions.

*STXBP1* encephalopathy shares the core clinical features with other synaptic vesicle cycle disorders, including intellectual disability, epilepsy, and motor dysfunctions. Thus, understanding the cellular and circuit origins of this disorder provides mechanistic insights into the growing list of neurodevelopmental disorders caused by presynaptic dysfunctions.

## Materials and Methods

### Mice

*Stxbp1* flox mice were generated from a previously described *Stxbp1* knockout-first allele (*tm1a*) that contains a trapping cassette flanked by two *FRT* sites and the exon 7 flanked by two *loxP* sites (Chen et al., 2020). *Stxbp1^tm1a/+^* mice were crossed to *Rosa26-Flpo* mice (JAX #012930)(Raymond and Soriano, 2007) to remove the trapping cassette in the germline, resulting in the *Stxbp1* flox allele (*tm1c*). *Stxbp1* flox mice were genotyped by PCR using a pair of primers 5’-TTCCACAGCCCTTTACAGAAAGG-3’ and 5’-ATGTGTATGCCTGGACTCACAGGG-3’ for both WT (352 bp) and *tm1c* (500 bp) alleles. *Stxbp1* flox mice were maintained on the C57BL/6J background by crossing to WT C57BL/6J mice (JAX # 000664). Heterozygous *Stxbp1* flox female mice (*Stxbp1^f/+^*) were crossed with C57BL/6J-congenic heterozygous *Vglut2-ires-Cre* (JAX # 028863)(Vong et al., 2011) male mice to generate *Stxbp1^f/+^;Vglut2^Cre/+^*, *Stxbp1^f/+^;Vglut2^+/+^*, *Stxbp^+/+^*, *Vglut2^cre/+^*, and WT mice. *Stxbp1^f/+^* female mice were also crossed with C57BL/6J-congenic heterozygous *Vgat-ires-Cre* (JAX # 028862)(Vong et al., 2011) male mice to generate *Stxbp1^f/+^;Vgat^Cre/+^*, *Stxbp1^f/+^*, *Vgat^+/+^*, *Stxbp1^+/+^*, *Vgat^Cre/+^*, and WT mice. Male white BALB/cAnNTac mice (Taconic # BALB-M) or BALB/cJ (JAX # 000651) were used for the resident-intruder test. Mice were housed in an Association for Assessment and Accreditation of Laboratory Animal Care International-certified animal facility on a 14-hour/10-hour light/dark cycle. All procedures to maintain and use mice were approved by the Institutional Animal Care and Use Committee at Baylor College of Medicine.

### Western blots

Western blot analyses were performed according to the protocols published previously (Chen et al., 2020) with minor modifications. Proteins were extracted from the entire mouse brains on postnatal day 0 or the cortices from 3-month-old mice. The lysis buffer contained 50 mM Tris-HCl (pH 7.6), 150 mM NaCl, 1 mM EDTA, 1% Triton X-100, 0.5% Na-deoxycholate, 0.1% SDS, and 1 tablet of cOmplete™, Mini, EDTA-free Protease Inhibitor Cocktail (Roche, catalog # SKU 11836170001) in 10 ml buffer. Stxbp1 was detected by a rabbit antibody against the N terminal residues 58–70 (Abcam, catalog # ab3451, lot #GR79394-18, 1:2,000 or 1:5,000 dilution) or a rabbit antibody against the C terminal residues 580–594 (Synaptic Systems, catalog # 116002, lot # 116002/15, 1:2,000 or 1:5,000 dilution). Gapdh was detected by a rabbit antibody (Santa Cruz Biotechnology, catalog # sc-25778, lot # A0515, 1:300 or 1:1,000 dilution). Primary antibodies were detected by a goat anti-rabbit antibody conjugated with IRDye 680LT (LI-COR Biosciences, catalog # 925-68021, lot # C40917-01, 1:20,000 dilution). Proteins were visualized and quantified using an Odyssey CLx Imager and Image Studio Lite version 5.0 (LI-COR Biosciences). Stxbp1 levels were normalized by the Gapdh levels. Each mouse was tested by both Stxbp1 antibodies and the results from the two antibodies were averaged.

### Double fluorescent *in situ* hybridization and imaging

A digoxigenin (DIG)-labeled RNA antisense probe against mouse *Stxbp1* and fluorescein (FITC)-labeled RNA antisense probes against mouse *Vglut1* (*Slc17a7*), *Vglut2* (*Slc17a6*), or *Gad1* were generated by *in vitro* transcription using cDNA templates and RNA DIG- or FITC-labeling kits (Sigma, catalog # 11277073910 or 11685619910, respectively). The DNA templates were made by PCR amplification from a plasmid pCMV-SPORT6-Stxbp1a (GenBank: BC031728.1, Transomic Technologies) for the *Stxbp1* probe or from mouse brain cDNA for the *Vglut1*, *Vglut2*, and *Gad1* probes, with a SP6 promoter (ATTTAGGTGACACTATAG) or a T3 promoter (AATTAACCCTCACTAAAGGG) added at the 5’ end of the PCR forward primers and a T7 promoter (TAATACGACTCACTATAGGG) at the 5’ end of the PCR reverse primers. The sequences of *Stxbp1*, *Vglut1*, and *Vglut2* probes were from Allen Brain Atlas (http://mouse.brain-map.org) and *Gad1* from Eurexpress (http://www.eurexpress.org/ee/). The probe sequences are listed in the ***Supplementary File 1***.

Double fluorescent *in situ* hybridization (DFISH) was performed by the RNA *In Situ* Hybridization Core at Baylor College of Medicine using an automated robotic platform and procedures as described previously (Yaylaoglu et al., 2005) with minor modifications for double ISH. Briefly, fresh-frozen brains were embedded in optimal cutting temperature (OCT) compound and cryosectioned (25 µm). Two or three probes were hybridized to brain sections simultaneously (*Stxbp1*/*Gad1* or *Stxbp1*/*Vglut1*/*Vglut2*) in hybridization buffer (Ambion, catalog # B8807G). Sections were washed with standard saline citrate stringency solution (SSC; 0.15 M NaCl, 0.015 M sodium citrate) to remove unbound and non-specifically bound probes. To visualize the DIG-labeled probe, brain sections were incubated for 30 minutes with a horse radish peroxidase (HRP)-conjugated sheep anti-DIG primary antibody (Sigma, catalog # 11207733910) diluted at 1/500 in Tris-NaCl blocking buffer (TNB; 100 mM Tris, 150 mM NaCl, 0.5% (w/v) blocking reagent (Perkin Elmer, catalog # FP1012), pH 7.6). After washes in Tris-NaCl-Tween (TNT; 10 mM Tris-HCl, pH 8.0, 150 mM NaCl and 0.05% TWEEN 20) buffer, brain sections were then developed with tyramide-Cy3 Plus (Akoya Biosciences, catalog # NEL744001KT, 1/50 dilution in amplification diluent, 15 minutes). After washes in TNT buffer, the remaining HRP activity was quenched by a 10-minute incubation in 0.2 M HCl. Sections were then washed in TNT, blocked in TNB for 15 minutes before incubation with an HRP-conjugated sheep anti-FITC antibody (Sigma, catalog # 11426346910) diluted at 1/500 in TNB for 30 minutes. After washes in TNT, the FITC-labeled probe was visualized using tyramide-FITC Plus (Akoya Biosciences, catalog # NEL741001KT, 1/50 dilution in amplification diluent, 15 minutes). The slides were washed in TNT and stained with 4’,6-diamidino-2-phenylindole (DAPI; Invitrogen, catalog # D3571), washed again, removed from the machine, and mounted in ProLong Diamond (Invitrogen, catalog # P36961).

Vglut2-cHet or Vgat-cHet and their respective control mice were processed, and brain sections were stained and imaged in parallel. Fluorescent images of brain sections were acquired using an Axio Zoom.V16 fluorescence microscope (Zeiss) and processed using Imaris (Oxford Instruments) or ImageJ (National Institutes of Health). The frontal cortex, somatosensory cortex, hippocampus, thalamus, reticular thalamic nucleus, striatum, and cerebellum were analyzed from sagittal sections, and amygdala and hypothalamus from coronal sections. 3–8 sections from each mouse were analyzed for each brain region. For *Vglut1/2*- or *Gad1*-positive cells, individual somas were selected using the surface function of Imaris with the following parameters: surface detail = 0.811, diameter of the largest square = 25 μm for cortical pyramidal neurons and Purkinje cells, and 20 μm for other neurons; pixels with the intensity at the lower 2–4% range of the maximal intensity were removed; voxels with the size at the lower 1.5–2% range of the maximal voxel size was removed. The mean intensity of *Stxbp1* was measured in each of the selected somas and then the average intensity was calculated across all selected cells for a brain section. For *Vglut1/2*- or *Gad1*-negative cells, *Vglut1/2*- or *Gad1*-positive cells were first selected as described above and removed. Individual *Vglut1/2*- or *Gad1*-negative somas were then selected based on *Stxbp1* signals using the parameters described above. The mean intensity of *Stxbp1* was measured in each of the selected somas and the average intensity was calculated across all selected cells for a brain section. Approximately 200–600 *Vglut1/2*-positive or *Gad1*-negative cells and 50–100 *Gad1*-positive or *Vglut1/2*-negative cells were selected for a brain region except the striatum where about 800 cells were selected in each section. For the hippocampal pyramidal neurons and cerebellar granular cells, due to their high cellular densities, the soma region of a group cells instead of individual cells was selected manually and the mean *Stxbp1* intensities were measured using ImageJ. Background signals were measured in intercellular space and subtracted from each measurement. *Stxbp1* levels from different brain sections were normalized by the average *Stxbp1* levels of WT brain sections that were simultaneously stained and imaged.

### Health monitoring

Body weight and dystonia of mice were monitored weekly. Dystonia was assessed by holding mice on their tails briefly in air and observing the movement of hindlimbs. Dystonia was scored as 0 = no dystonia, 1 = stiffness in hindlimb, 2 = clasping of one hindlimb, 3 = clasping of both hindlimbs, 4 = tight clasping of both hindlimbs.

### Behavioral tests

All behavioral experiments were performed using the equipment and facility at the Neurobehavioral Core of Baylor College of Medicine Intellectual and Developmental Disabilities Research Center. All tests were performed and analyzed blind to the genotypes according to the protocols published previously (Chen et al., 2020) with minor modifications. Approximately equal numbers of cHet mice and their sex- and age-matched WT, Flox and Cre littermates were tested in parallel in each experiment except for resident intruder test where only male mice were used. Mice were habituated in the behavioral test facility for at least 30 minutes before testing. The sexes and ages of the tested mice were indicated in the figures.

Nesting test: An autoclaved Nestlet was given to a mouse individually housed in its home cage, and the quality of the nest was scored after 24 hours.

Elevated plus maze test: A mouse was placed in the center of an elevated maze consisting of two open arms (25 × 8 cm) and two closed arms with high walls (25 × 8 × 15 cm). The mouse was initially placed facing the open arms and then allowed to freely explore for 10 minutes with 150–200-lux illumination and 65-dB background white noise. The mouse activity was recorded using a video camera (ANY-maze, Stoelting).

Open-field test: A mouse was placed at the center of a clear, open chamber (40 × 40 × 30 cm) and allowed to freely explore for 30 minutes with 150–200-lux illumination and 65-dB background white noise. The horizontal plane was evenly divided into 256 squares (16 × 16), and the center zone is defined as the central 100 squares (10 × 10). The horizontal travel and vertical activity were quantified by either an Open Field Locomotor system or a VersaMax system (OmniTech).

Marble burying test: A clean standard housing cage was filled with approximately 8-cm deep bedding material. 20 marbles were arranged on top of the bedding in a 4 × 5 array. A mouse was placed into this cage for 30 minutes before the number of buried marbles (i.e., at least 50% of the marble covered by the bedding material) was recorded.

Rotarod test: A mouse was tested on an accelerating rotarod apparatus (Ugo Basile) in 3 trials per day for 4 consecutive days. There was a 30–60-minute resting interval between trials. Each trial lasted for a maximum of 5 minutes, during which the rod accelerated linearly from 4 to 40 revolutions per minute (RPM). The time when the mouse walks on the rod and the latency for the mouse to fall from the rod were recorded for each trial.

Foot slip test: A mouse was placed onto an elevated 40 × 25 cm wire grid (1 × 1 cm spacing) and allowed to freely move for 5 minutes. The number of foot slips was manually counted, and the moving distance was measured through a video camera (ANY-maze, Stoelting). The number of foot slips were normalized by the moving distance for each mouse.

Vertical pole test: A mouse was placed at the top of a vertical threaded metal pole (1.3-cm diameter, 55-cm length). The amount of time for the mouse to descend to the floor was measured with a maximal cutoff time of 120 seconds.

Acoustic startle response test: A mouse was placed in a plastic cylinder and acclimated to the 70-dB background white noise for 5 minutes. The mouse was then tested with 4 blocks, and one block consisted of 13 trials. In one block, each of 13 different levels of sound (70, 74, 78, 82, 86, 90, 94, 98, 102, 106, 110, 114, or 118 dB, 40 ms, inter-trial interval of 15 seconds on average) was presented in a pseudorandom order. The startle response was recorded for 40 ms after the onset of the sound. The rapid force changes due to the startles were measured by an accelerometer (SR-LAB, San Diego Instruments).

Pre-pulse inhibition test: A mouse was placed in a plastic cylinder and acclimated to the 70 dB background noise for 5 minutes. The mouse was then tested with 6 blocks, and one block consisted of 8 trials in a pseudorandom order: a “no stimulus” trial (40 ms, only 70 dB background noise present), a test pulse trial (40 ms, 120 dB), 3 different pre-pulse trials (20 ms, 74, 78, or 82 dB), and 3 different pre-pulse inhibition trials (a 20 ms, 74, 78, or 82 dB pre-pulse preceding a 40 ms, 120 dB test pulse by 100 ms). The startle response was recorded for 40 ms after the onset of the 120 dB test pulse. The inter-trial interval is 15 s on average. The rapid force changes due to the startles were measured by an accelerometer (SR-LAB, San Diego Instruments). Pre-pulse inhibition of the startle responses was calculated as “1 – (pre-pulse inhibition trial/test pulse trial)”.

Hot plate test: A mouse was placed on a hot plate (Columbus Instruments) with a temperature of 55 °C. The latency for the mouse to first respond with either a hind paw lick, hind paw flick, or jump was recorded. If the mouse did not respond within 45 seconds, then the test was terminated, and the latency was recorded as 45 seconds.

Three-chamber test: The apparatus (60.8 × 40.5 × 23 cm) consists of three chambers (left, center, and right) of equal size with 10 × 5 cm openings between the chambers. A test mouse was placed in the apparatus with a mesh pencil cup in each of the left and right chambers and allowed to freely explore for 10 minutes. A novel object was then placed under one mesh pencil cup and an age- and sex-matched partner mouse (WT C57BL/6J) under the other mesh pencil cup. The test mouse was allowed to freely explore for another 10 minutes. The position of the test mouse was tracked through a video camera (ANY-maze, Stoelting), and the approaches of the test mouse to the object or partner mouse were scored manually. Partner mice were habituated to the mesh pencil cups in the apparatus for 1 hour per day for 2 days prior to testing. A partner mouse was used only in one test per day.

Resident-intruder test: Male test mice (resident mice) were individually caged for 2 weeks before testing. Age-matched male BALB/cAnNTac or BALB/cJ mice were group-housed to serve as the intruders. During the test, an intruder was placed into the home cage of a test mouse for 10 minutes, and their behaviors were video recorded. Videos were scored for the number and duration of each attack by the resident mouse regardless the attack was initiated by either the resident or intruder.

Novel object recognition test: A mouse was first habituated in an empty arena (24 × 45 × 20 cm) for 5 minutes before every trial, and then placed into the testing arena with two identical objects (i.e., familiar object 1 and familiar object 2) for the first three trials. In the fourth trial, familiar object 1 was replaced with a novel object. In the fifth trial, the mouse was presented with the two original, identical objects again. Each trial lasted 5 minutes. The inter-trial interval was 24 hours.

The movement of mice was recorded by a video camera. The amount of time that the mouse interacted with the objects (*T*) was recorded using a wireless keyboard (ANY-maze, Stoelting). The preference index of interaction was calculated as *T_familiar object 1_ /(T_familiar object 1_ + T_familiar object 2_)* for the first three trials and fifth trial and as *T_novel object_/(T_novel object_ + T_familiar object 2_)* for the fourth trial.

Fear conditioning test: Pavlovian fear conditioning was conducted in a chamber (30 × 25 × 29 cm) with a grid floor for delivering electrical shocks (Coulbourn Instruments). During the 5-minute training phase, a mouse was placed in the chamber for 2 minutes, and then a sound (85 dB, white noise) was turned on for 30 seconds immediately followed by a mild foot shock (2 sec, 0.72 mA). The same sound and foot shock were repeated one more time 2 minutes after the first foot shock. After the second foot shock, the mouse stayed in the training chamber for 18 seconds before returning to its home cage. After 24 hours, the mouse was tested for the contextual and cued fear memories. In the contextual fear test, the mouse was placed in the same training chamber and its freezing behavior was monitored for 5 minutes without the sound stimulus. The mouse was then returned to its home cage. One to two hours later, the mouse was transferred to the chamber after it has been altered using plexiglass inserts and a different odor to create a new context for the cued fear test. After 3 minutes in the chamber, the same sound cue that was used in the training phase was turned on for 3 minutes without foot shocks while the freezing behavior was monitored. The freezing behavior was scored using an automated video-based system (FreezeFrame, Actimetrics). The freezing time (%) during the first 2 minutes of the training phase (i.e., before the first sound) was subtracted from the freezing time (%) during the contextual fear test. The freezing time (%) during the first 3 minutes of the cued fear test (i.e., without sound) was subtracted from the freezing time (%) during the last 3 minutes of the cued fear test (i.e., with sound).

### Video-EEG/EMG

Video-EEG/EMG recordings and analysis were performed as previously described (Chen et al., 2020). Briefly, mice at 8–13 weeks of age were anesthetized with 2.5% isoflurane in oxygen. Approximately 0.25 mm-diameter craniotomies were performed at the coordinates below that were normalized by the distance between bregma and lambda (DBL). Perfluoroalkoxy polymer (PFA)-coated silver wire electrodes (A-M Systems, catalog # 786000, 127 mm bare diameter, 177.8 mm coated diameter) were used for grounding at the right frontal cortex, referencing at the cerebellum, and recording at the left frontal cortex (anterior posterior (AP): 0.42 of DBL, medial lateral (ML): 0.356 of DBL, dorsal ventral (DV): −1.5 mm), left, and right somatosensory cortices (AP: −0.34 of DBL, ML: ± 0.653 of DBL, DV: −1.5mm). An EMG recording and an EMG reference electrode were inserted into the neck muscles. The mice were allowed to recover from the surgeries for at least one week. Before recording, mice were individually habituated in the recording chambers for 24 hours. EEG/EMG signals (5000-Hz sampling rate with a 0.5-Hz high-pass filter) and videos (30 frames per second) were recorded synchronously for continuous 72 hours using a 4-channel EEG/EMG tethered system (Pinnacle Technology).

Spike-wave discharges (SWDs) were identified by generating putative candidates with custom written code in MATLAB (MathWorks) followed by the classification of candidates with a convolutional neural network in Python that has been trained with manually labeled EEG segments (Chen et al., 2020). Myoclonic seizures were identified by visual inspection of EEG/EMG signals and videos to identify sudden jumps and jerks (Chen et al., 2020). The state of the mouse before each myoclonic seizure event was classified as REM sleep, NREM sleep, or awake based on the EEG/EMG. The video component of the data file for one Vglut2-cHet mouse was corrupted, precluding the identification of myoclonic seizures. Thus, this mouse was only analyzed for SWDs.

### Statistics

All reported sample numbers (*n*) represent independent biological replicates that are the numbers of tested mice or tissue sections. Statistical analyses were performed with Prism 8 or 9 (GraphPad Software) unless stated otherwise below. Anderson-Darling test, D’Agostino-Pearson, Shapiro-Wilk, and Kolmogorov-Smirnov tests were used to determine if data were normally distributed. If all data within one experiment passed all four normality tests, then the statistical test that assumes a Gaussian distribution was used. Otherwise, the statistical test that assumes a non-Gaussian distribution was used. Nested one-way ANOVA with Turkey multiple comparison was used to assess statistical significance of *Stxbp1* expression levels in the DFISH experiments. Either one-way or two-way ANOVA with multiple comparisons was used for Western blot, behavior, and EEG data analyses. For data with Gaussian distribution, ordinary one-way ANOVA with Turkey multiple comparison was used. For non-Gaussian distributed data, Kruskal-Wallis one-way ANOVA with Dunn’s multiple comparison test was used. Turkey multiple comparison test was also used in conjunction with two-way ANOVA. For body weight, dystonia score, rotarod test, acoustic startle response, and novel object recognition, two-way repeated measures ANOVA was used with Turkey multiple comparison. Either two-way or three-way ANOVA with Turkey multiple analysis was used for gender effect analyses.

OriginPro 2021 (OriginLab) was used to perform three-way ANOVA. To compare the difference between the dystonia scores of Vglut2-cHet and Vgat-cHet mice, Fisher-Freeman-Halton exact test was performed using StatXact 12 (Cytel). The details of all statistical tests, numbers of replicates, and *P* values are reported in **Supplementary File 3**.

## Acknowledgments

We thank the RNA In Situ Hybridization Core at Baylor College of Medicine for performing the DFISH experiments with the expert assistance of Dr. Cecilia Ljungberg. This work was supported in part by Citizens United for Research in Epilepsy (CURE Epilepsy Award to MX), the National Institute of Neurological Disorders and Stroke (R01NS100893 to MX), the National Institute of Mental Health (R01MH117089 to MX), the Eunice Kennedy Shriver National Institute of Child Health and Human Development (P50HD103555 to Baylor College of Medicine Intellectual and Developmental Disabilities Research Center, Neurovisualization Core and Neurobehavioral Core), and the National Institutes of Health (S10OD016167 to Dr. Cecilia Ljungberg). MX is a Caroline DeLuca Scholar. This article is dedicated to the memory of Caroline DeLuca, who inspired this project.

---

**Figure 1-supplement 1.**
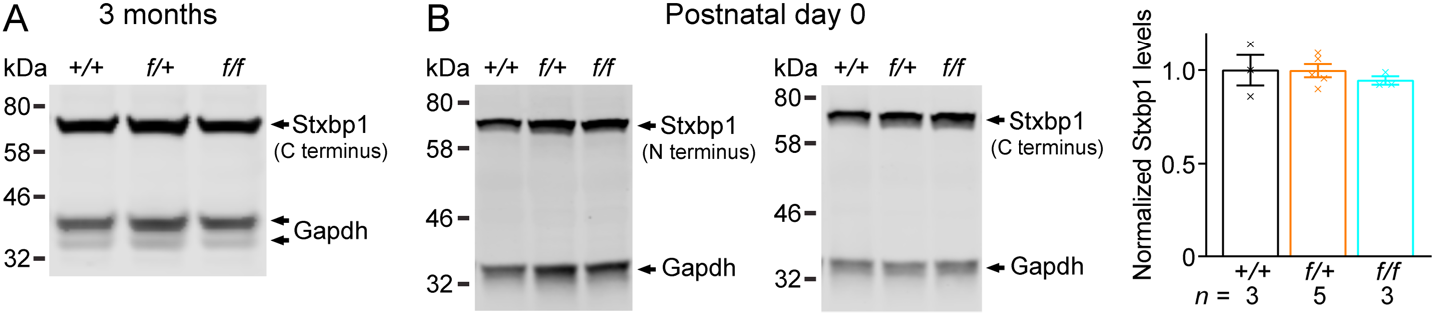
Normal Stxbp1 protein levels in *Stxbp1^f/+^ and Stxbp1^f/f^* mice. (**A**) A representative Western blot of proteins from the cortices of 3-month-old WT, *Stxbp1^f/+^*, and *Stxbp1^f/f^* mice. Stxbp1 was detected by an antibody recognizing its C terminus. Gapdh, a housekeeping protein as loading control. (**B**) Left, representative Western blots of proteins from the brains of WT, *Stxbp1^f/+^*, and *Stxbp1^f/f^* mice at postnatal day 0. Stxbp1 was detected by an antibody recognizing its N terminus (left blot) or C terminus (right blot). Right, summary data of normalized Stxbp1 protein levels at postnatal day 0. Stxbp1 levels were first normalized by the Gapdh levels and then by the average Stxbp1 levels of all WT mice from the same blot. Each cross represents one mouse. Bar graphs are mean ± s.e.m.

**Figure 1-supplement 2.**
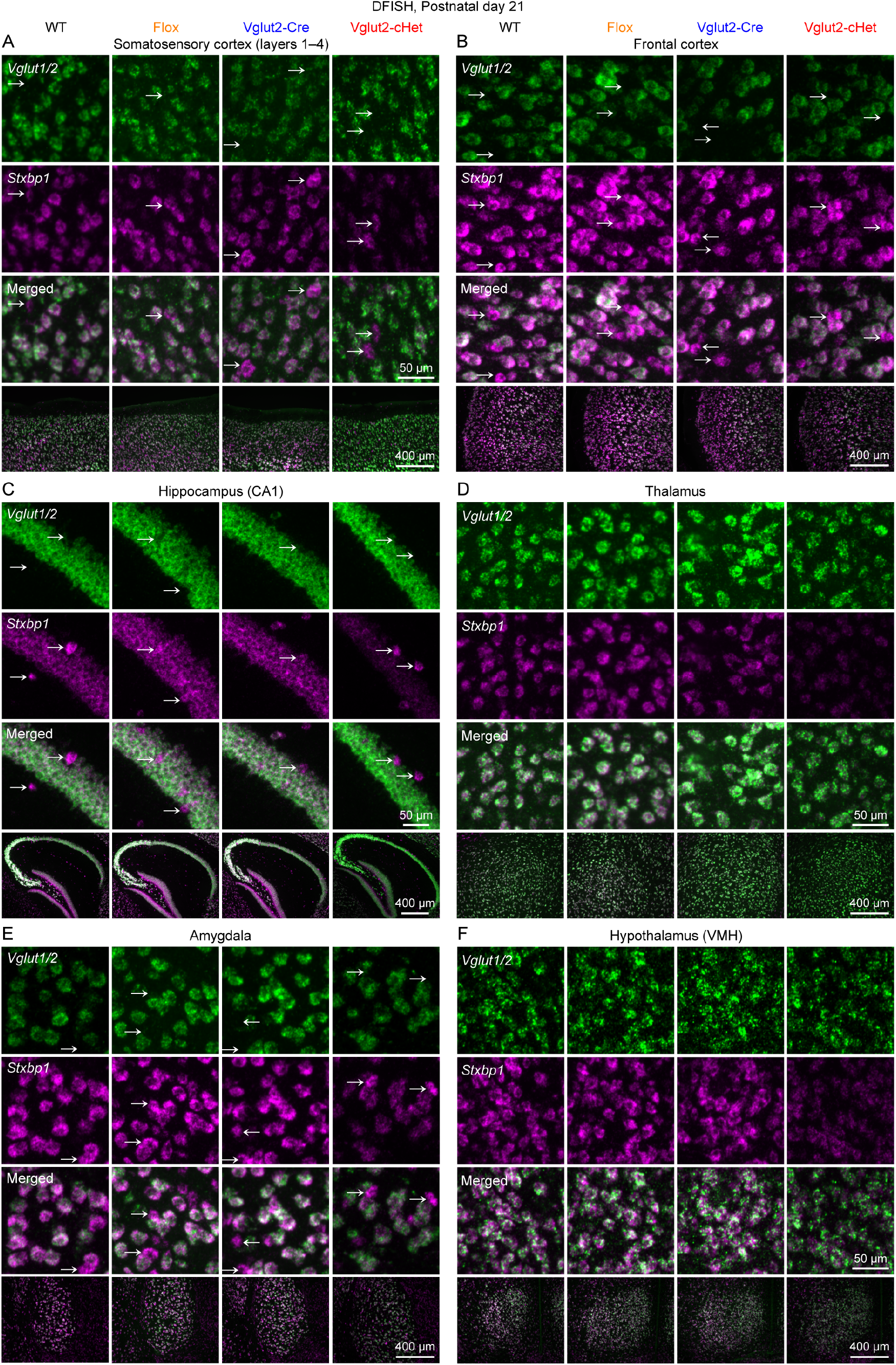
Reduction of *Stxbp1* levels specifically in glutamatergic neurons of Vglut2-cHet mice (Part 1). (**A**) Representative fluorescent images from brain sections labeled by ISH probes against *Stxbp1* and *Vglut1/2*. The bottom row shows the layers 1–4 of the somatosensory cortices, and the top three rows show the individual cells from this region. Arrows indicate *Vglut1/2*-negative cells. (**B–F**) Similar to (A), but for other brain regions indicated on the top of each panel. VMH, ventromedial hypothalamic nucleus.

**Figure 1-supplement 3.**
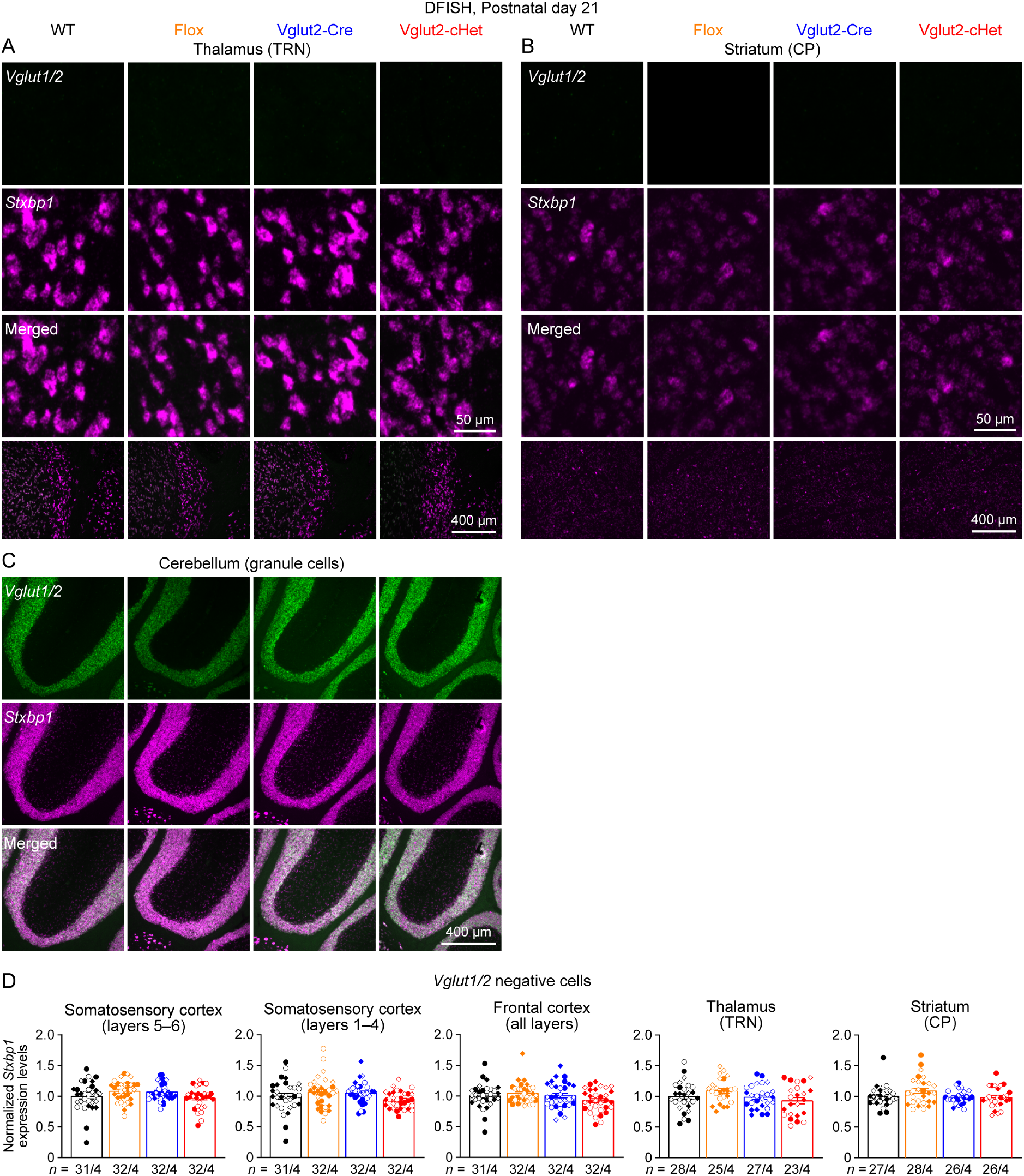
Reduction of *Stxbp1* levels specifically in glutamatergic neurons of Vglut2-cHet mice (Part 2). (**A**) Representative fluorescent images from brain sections labeled by ISH probes against *Stxbp1* and *Vglut1/2*. The bottom row shows the thalamic reticular nucleus (TRN), and the top three rows show the individual cells from this region. (**B,C**) Similar to (A), but for other brain regions indicated on the top of each panel. (**D**) Summary data of normalized *Stxbp1* mRNA levels in *Vglut1/2*-negative cells from different brain regions. *Stxbp1* levels were normalized by the average *Stxbp1* levels of WT brain sections that were simultaneously stained and imaged. The *Stxbp1* levels of Vglut2-cHet mice were normal. Different shapes of symbols represent different mice (4 mice per genotype, filled circle and diamond for 2 males and open circle and diamond for 2 females), and each symbol represents one brain section. TRN, thalamic reticular nucleus; CP, caudoputamen. Bar graphs are mean ± s.e.m.

**Figure 1-supplement 4.**
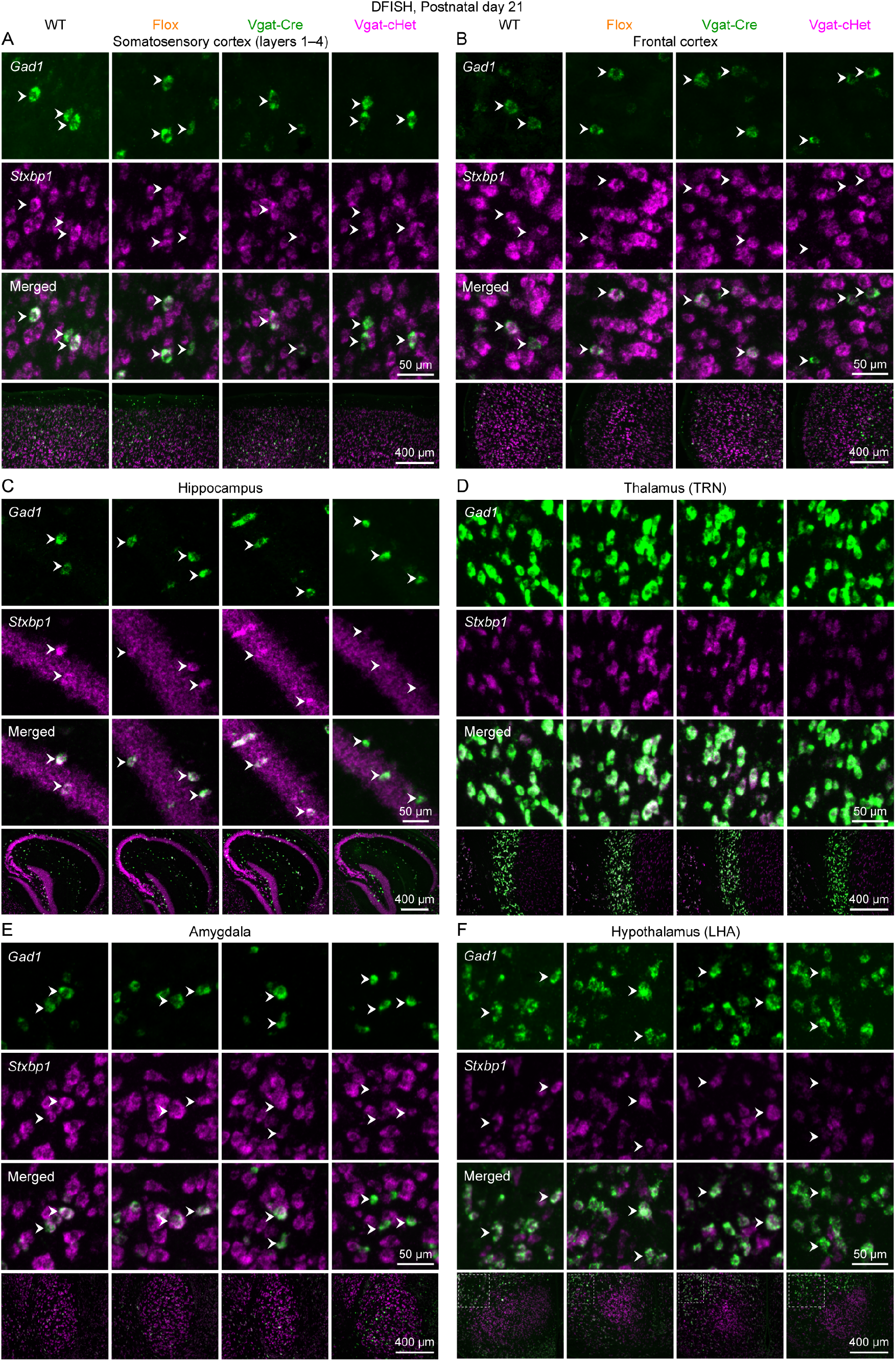
Reduction of *Stxbp1* levels specifically in GABAergic neurons of Vgat-cHet mice (Part 1). (**A**) Representative fluorescent images from brain sections labeled by ISH probes against *Stxbp1* and *Gad1*. The bottom row shows the layers 1–4 of the somatosensory cortices, and the top three rows show the individual cells from this region. Arrow heads indicate *Gad1*-positive cells. (**B–F**) Similar to (A), but for other brain regions indicated on the top of each panel. TRN, thalamic reticular nucleus; LHA, lateral hypothalamic area.

**Figure 1-supplement 5.**
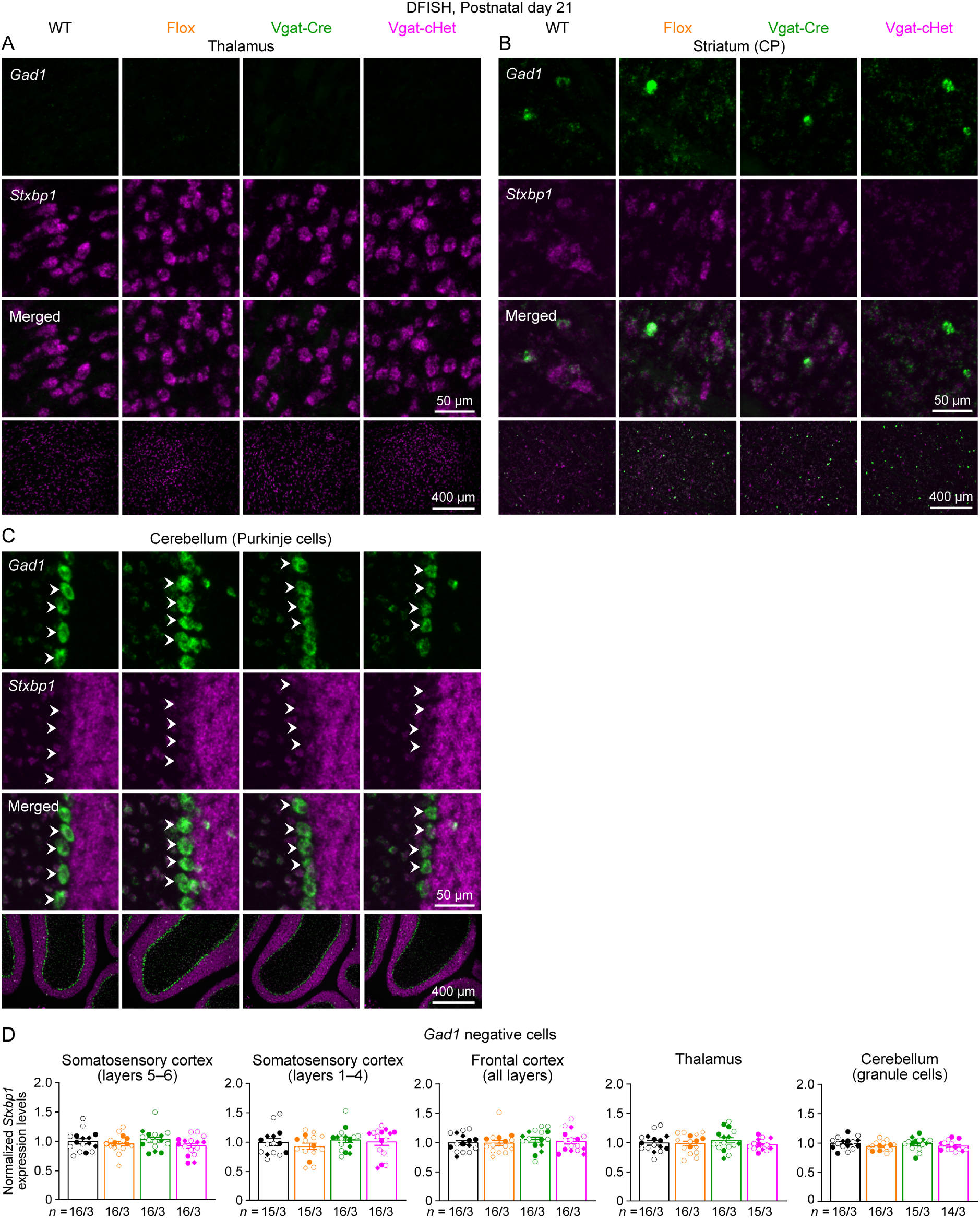
Reduction of *Stxbp1* levels specifically in GABAergic neurons of Vgat-cHet mice (Part 2). (**A**) Representative fluorescent images from brain sections labeled by ISH probes against *Stxbp1* and *Gad1*. The bottom row shows the thalamus, and the top three rows show the individual cells from this region. (**B,C**) Similar to (A), but for other brain regions indicated on the top of each panel. (**D**) Summary data of normalized *Stxbp1* mRNA levels in *Gad1*-negative cells from different brain regions. *Stxbp1* levels were normalized by the average *Stxbp1* levels of WT brain sections that were simultaneously stained and imaged. The *Stxbp1* levels of Vgat-cHet mice were normal. Different shapes of symbols represent different mice (3 mice per genotype, filled circle and diamond for males and open circle and diamond for females), and each symbol represents one brain section. CP, caudoputamen. Bar graphs are mean ± s.e.m.

**Figure 3-supplement 1.**
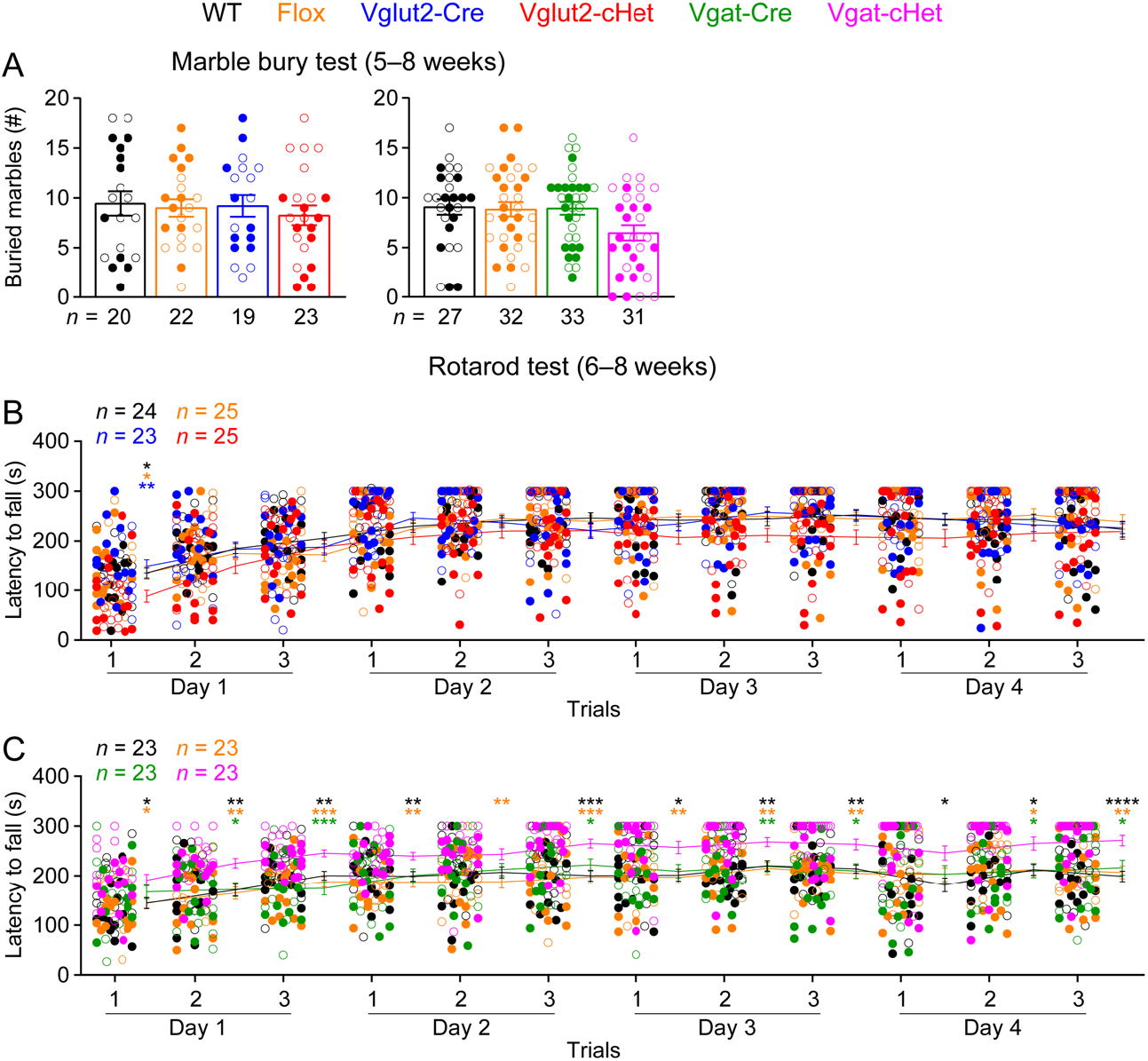
Vglut2-cHet and Vgat-cHet mice do not show impairments in marble bury and rotarod tests. (**A**) The numbers of buried marbles by Vglut2-cHet and Vgat-cHet mice were not significantly different from those of control mice. (**B,C**) In the rotarod test, Vglut2-cHet mice performed similar to control mice except the first trial (B) and Vgat-cHet mice performed better than control mice, as they were able to stay on the rotating rod for longer time (C), probably due to their lower body weight or hyperactivity. The statistical significance between Vglut2-cHet and WT, Flox, or Vglut2-Cre mice is indicated by black, orange, or blue asterisks, respectively, and between Vgat-cHet and WT, Flox, or Vgat-Cre mice by black, orange, or green asterisks, respectively. The numbers and ages of tested mice are indicated in the figures. Each filled (male) or open (female) circle represents one mouse. Bar graphs are mean ± s.e.m. * *P* < 0.05, ** *P* < 0.01, *** *P* < 0.001, **** *P* < 0.0001.

**Figure 3-supplement 2.**
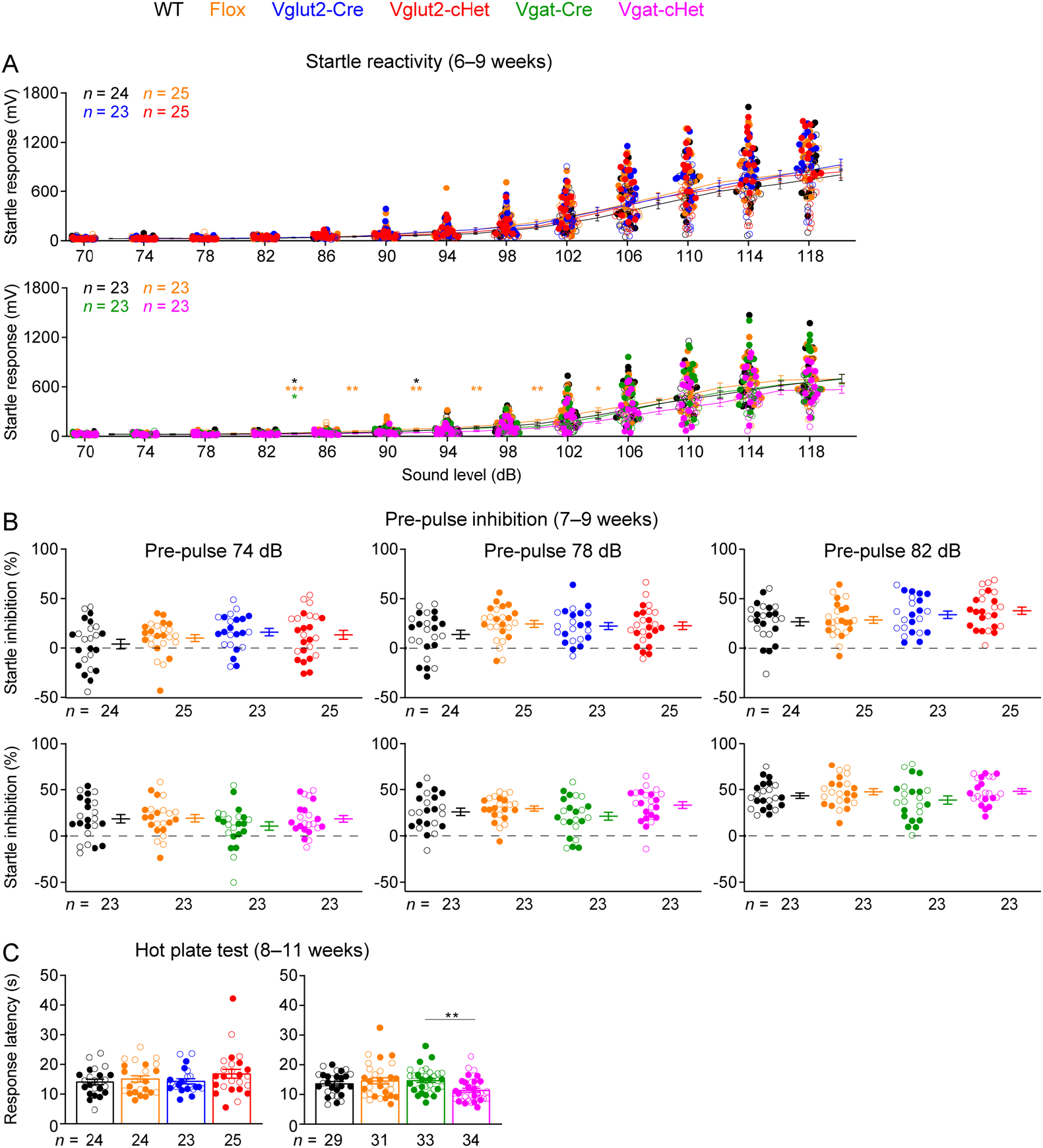
Vglut2-cHet and Vgat-cHet mice have normal sensory functions. (**A**) Vglut2-cHet and Vgat-cHet mice showed similar acoustic startle responses as control mice at different sound levels. The statistical significance between Vgat-cHet and WT, Flox, or Vgat-Cre mice is indicated by black, orange, or green asterisks, respectively. (**B**) In the pre-pulse inhibition test, when a weak sound (i.e., pre-pulse 74, 78, or 82 dB) preceded a loud sound (120 dB), Vglut2-cHet and Vgat-cHet mice showed a similar reduction in the startle responses to the loud sound as control mice. (**C**) In the hot plate test, Vglut2-cHet mice showed similar latencies in response to the high temperature as control mice, and the latencies of Vgat-cHet were slightly shorter than Vgat-Cre mice. The numbers and ages of tested mice are indicated in the figures. Each filled (male) or open (female) circle represents one mouse. Bar graphs are mean ± s.e.m. * *P* < 0.05, ** *P* < 0.01, *** *P* < 0.001.

**Figure 5-supplement 1.**
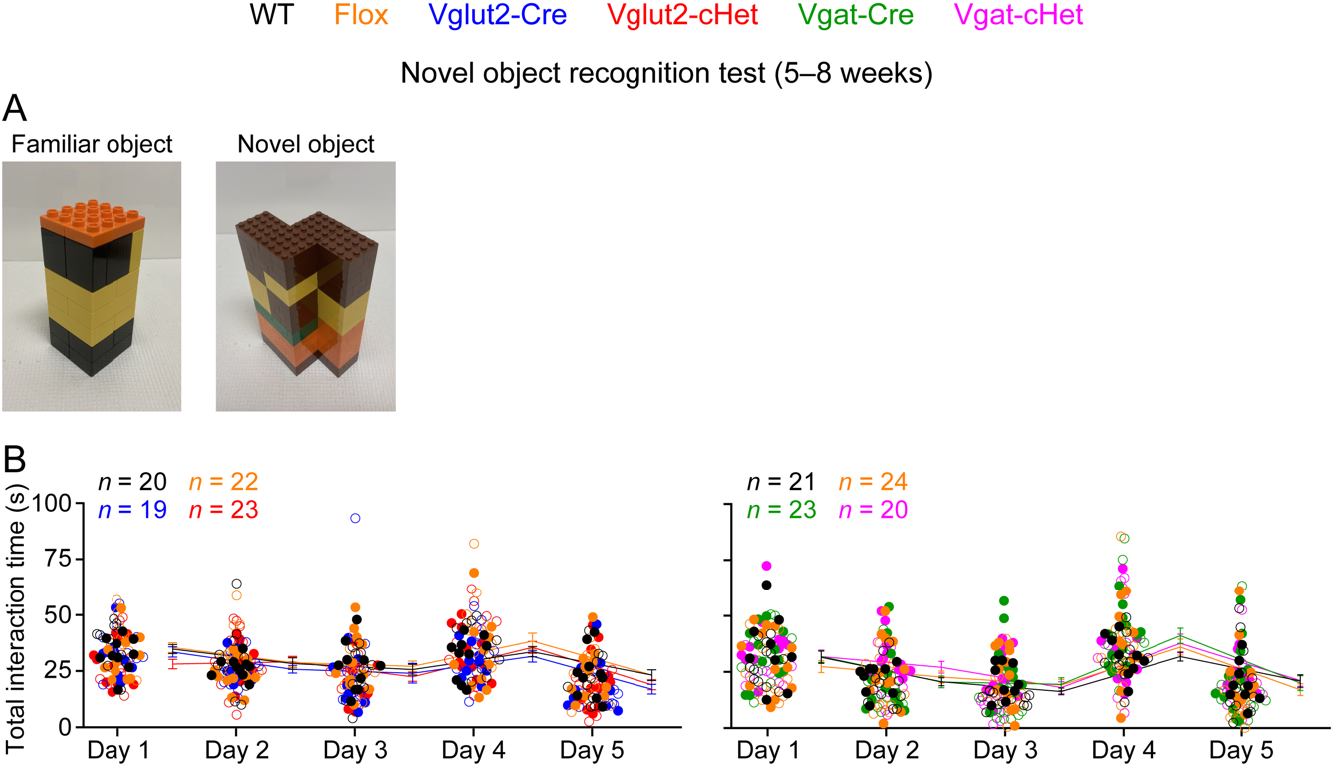
Normal novel objection recognition in Vglut2-cHet and Vgat-cHet mice. (**A**) The images of the familiar and novel objects used in the novel objection recognition test. (**B**) The total interaction time with familiar and novel objects of Vglut2-cHet or Vgat-cHet mice was similar to that of control mice. The numbers and ages of tested mice are indicated in the figures. Each filled (male) or open (female) circle represents one mouse. Bar graphs are mean ± s.e.m.

**Supplementary File 1. Double fluorescent *in situ* hybridization probe sequences**

The sequences of the *Stxbp1*, *Vglut1*, *Vglut2*, and *Gad1* probes are provided.

**Supplementary File 2. Phenotypic comparison of human patients and different mouse models.**

The phenotyping tests in different mouse models (the second column) are grouped based on the clinical features of *STXBP1* encephalopathy (the first column). The results of the phenotyping tests from different mouse models are compared in the table.

**Supplementary File 3. Statistics of experimental results.**

The details of all statistical tests, numbers of replicates, and *P* values are presented for each experiment in the table.

**Video 1. Vglut2-cHet mice show SWDs.**

A representative video showing several SWDs in a Vglut2-cHet mouse. The top 3 traces are EEG signals from the left frontal cortex, right somatosensory cortex, and left somatosensory cortex. The bottom trace is the EMG signal from the neck muscle. The vertical line indicates the time of the current video frame. Note that the EEG signal from the left somatosensory cortex (the third channel) is inverted.

**Video 2. Vgat-cHet mice show myoclonic jerks.**

A representative video showing a myoclonic jerk of a Vgat-cHet mouse. The top 3 traces are EEG signals from the left frontal cortex, right somatosensory cortex, and left somatosensory cortex. The bottom trace is the EMG signal from the neck muscle. The vertical line indicates the time of the current video frame. The mouse was in REM sleep before the jerk. Note that the EEG signal from the left somatosensory cortex (the third channel) is inverted.

**Video 3. Vgat-cHet mice show myoclonic jumps.**

A representative video showing a myoclonic jump of a Vgat-cHet mouse. The top 3 traces are EEG signals from the left frontal cortex, right somatosensory cortex, and left somatosensory cortex. The bottom trace is the EMG signal from the neck muscle. The vertical line indicates the time of the current video frame. The mouse was in REM sleep before the jump. Note that the EEG signal from the left somatosensory cortex (the third channel) is inverted.

