## Supplementary File 1 for "Glutamatergic and GABAergic neurons mediate distinct neurodevelopmental phenotypes of *STXBP1* encephalopathy"

*Stxbp1* probe sense sequence (Allen Brain Atlas probe RP_040227_01_08):

GCGTCCTTCAGCACCACTGCTGTGAGTGCCCGCTATGGACATTGGCACAAGAATAAGGCCCCCGGGGAGTACCGCAGCGGTCCCCGCCTCATTATTTTCATCCTTGGGGGTGTGAGCCTGAATGAGATGCGCTGTGCTTACGAAGTGACCCAGGCCAACGGCAAGTGGGAAGTGCTGATAGGTTCTACTCACATTCTCACTCCCACCAAATTCCTCATGGACCTGAGACACCCCGACTTCAGGGAGTCCTCTAGGGTATCTTTTGAGGATCAGGCTCCAACAATGGAGTGAGAGCCAAAGAGACAAAGATCCACGCACATTCTCACCCCACAGAAACTGCTGGACACGCTGAAGAAGCTGAATAAAACAGATGAAGAAATAAGCAGTTAAAAAATAAGCTGCCCCCCAAAACCCCGGCTCCCTTCCCAAAATGCTCTGCAGCTCCCCCGTGCGCCACCTCGGTTACTCTGCTGCCTCCCCAGCCCTGCACGCCCTGGCCACCCCGTTGCCGTGCTGAGTTCTTCTCCTGTGCGATGACACCCCATCTTGTCCTCTGAAAAGCAAGAGAGTAATGTGTTGTTTTTTAAAAATGAGCATCTTCTGTATGTATCCCACAGTAAGTTCACATGCAAGCTCCACACTGCAGAAGCGTCAGAACTCCGGACCGAGTGAATTCTCCCTTATTTATGACCCCGTGACCTGTATATAGCCCTGTCCCGCGTGTGCACATTGCTTGAATATGGAAAGGTAGATGTGTGGGTGTCTCTCCAAGCTTGGTTGGATTCATTTCTGTCCTTGTTGGTGTTTGTTCCCCGGATAGGACATGCT

*Vglut1* probe sense sequence (Allen Brain Atlas probe RP_050310_01_B09):

CAGAGCCGGAGGAGATGAGCGAGGAGAAGTGTGGCTTTGTTGGCCACGACCAGCTGGCTGGCAGTGACGAAAGTGAAATGGAGGACGAGGCTGAGCCCCCAGGGGCGCCCCCCGCGCCGCCTCCGTCCTACGGGGCCACACACAGCACAGTGCAGCCTCCGAGGCCCCCGCCCCCTGTCCGGGACTACTGACCACGGGCCTCCCACTGTGGGGCAGTTTCCAGGACTTCCACTCCATACACCTCTAGCCTGAGCGGCAGTGTCGAGGAACCCCACTCCTCCCCTGCCTCAGGCTTAAGATGCAAGTCCTCCCTTGTTCCCAGTGCTGTCCGACCAGCCCTCTTTCCCTCTCAACTGCCTCCTGCGGGGGGTGAAGCTGCACACTAGCAGTTTCAAGGATACCCAGACTCCCCTGAAAGTCGTTCTCCGCTTGTTTCTGCCTGTGTGGGCTCAAATCTCCCCTTTGAGGGCTTTATTTGGAGGGACAGTTCAACCTCTTCCTCTCTTGTGGTTTTGAGGTTTCACCCCTTCCCCCAAGACCCCAGGGATTCTCAGGCTACCCCGAGATTATTCAGGTGGTCCCCTACTCAGAAGACTTCATGGTCGTCCTCTATTAGTTTCAAGGCTCGCCTAACCAATTCTACATTTTTCCAAGCTGGTTTAACCTAACCACCAATGCCGCCGTTCCCAGGACTGATTCTCACCAGCGTTTCTGAGGGA

*Vglut2* probe sequence (Allen Brain Atlas probe RP_050921_01_E03):

CCAAATCTTACGGTGCTACCTCACAGGAGAATGGAGGCTGGCCTAACGGCTGGGAGAAAAAGGAAGAATTTGTGCAAGAAGGTGCGCAAGACGCGTACACCTATAAGGACCGAGATGATTATTCATAACGATGCTAGTTGCTGGATTCATTTGTAGTGTTTGTGAATCAATTAATTGTGATTGCACAAAAATAATTTTAAAAATGTGGTGTGAACATGTAAACATATCAACCAAGCAAGTCTTGCTGTTCAAAAACAAAAACAAAAAAATCTGAATTCAAAACAGACCATGAGATTCCCATCAAGTGCAATCTGTGGCAGTTGTCACGTTATGCCGTCTTCATTCAGGCCATTTGTCCTTTCGTTTGTGATTTAAAGGTTTCCTGTAGAAATAAGTAGGTATTCGTTGGACCCATCACCATTTTAGAGAGCACAACTACAACAGTTGGCACATGTCATCCTACAGAAGTTAGGAAGCCAAAGCTACTGGATCATGCAAACTGCACTTATTTATTACACTGGACTGCAAACTATCCCAGGGAAAGCCTGTCTAGAGACATAGTGGAACAGGAAAGATGGCT

*Gad1* probe sequence (Eurexpress probe T3752)

TGGCCTCGAGCCAGATTCGGACGAGGACCAGGGATCGTGCAAGCAAGGAAGCAGCCCTGGGGTGACACCCAGCACGTACTCCTGTGACAGAGCCGAGCCCAGCCCAGCCCCGGGACGCTTCGCAGAGGAGTCGCGGGAGGGTCCAGCTCGCTGTCGCTGAACCGAGCCTGTTCCTGCGCCCAGTCTGCGGGGGACCCTTGAACCGTAGAGACCCCAAGACCACCGAGCTGATGGCATCTTCCACTCCTTCGCCTGCAACCTCCTCGAACGCGGGAGCGGATCCTAATACTACCAACCTGCGCCCTACAACGTATGATACTTGGTGTGGCGTAGCCCATGGATGCACCAGAAAACTGGGCCTGAAGATCTGTGGCTTCTTACAAAGGACCAATAGCCTGGAAGAGAAGAGTCGTCTTGTGAGCGCCTTCAGGGAGAGGCAGTCCTCCAAGAACCTGCTTTCCTGTGAAAACAGTGACCAGGGTGCCCGCTTCCGGCGCACAGAGACCGACTTCTCCAACCTGTTTGCTCAAGATCTGCTTCCAGCTAAGA
