## Supplementary File 2 for "Glutamatergic and GABAergic neurons mediate distinct neurodevelopmental phenotypes of *STXBP1* encephalopathy"

**Table S1. Phenotypic comparison of *Stxbp1* constitutive and cell-type specific haploinsufficiency mouse models**

| Human patient phenotypes  (% of patients^1^) | Mouse phenotyping tests | Mouse *Stxbp1* haploinsufficiency models (C57BL/6J)  and phenotypes^2^ | | |
| --- | --- | --- | --- | --- |
|  |  | *Stxbp1^tm1d/+^* | Vglut2-cHet | Vgat-cHet |
|  |  | Chen et al., 2020 | This paper | This paper |
| – | Stxbp1 protein or mRNA reduction in brain tissues | Protein:  40–50% in cortex, hippocampus, thalamus + hypothalamus, striatum, midbrain + hindbrain;  30% in cerebellum;  20% in olfactory bulb | mRNA:  50-60% reduction in frontal cortex, thalamus, amygdala, hypothalamus (VMH);  40% in hippocampus, somatosensory cortex;  17% in cerebellar granular cells | mRNA:  50-60% reduction in frontal cortex, somatosensory cortex, thalamic reticular nucleus, striatum; 40% in hippocampus, hypothalamus (LHA), amygdala, cerebellar Purkinje cells |
| Epilepsy (95%) | Video-EEG/EMG | SWDs and myoclonic seizures | SWDs | Myoclonic seizures |
| Intellectual disability (100%) | Novel object recognition | Yes | No | No |
|  | Contextual fear | Yes | No | Yes |
|  | Cued fear | Yes | Yes | No |
| Motor deficits  (92%) | Hindlimb clasping | Yes | No^3^ | Yes |
|  | Foot slip | Yes | No | Yes |
|  | Vertical pole | Yes | No | Yes |
|  | Rotarod | No^4^ | No | No^4^ |
| Developmental delay (64.3%) | Body weight | Yes | No | Yes |
| Hyperactivity (4%) | Open-field | Yes | No | Yes |
| Autistic traits (17%) | Three-chamber | No | No | No |
| Aggressive behavior (3.4%) | Resident- intruder | Yes | No | Yes |
| Anxiety  (27%) | Elevated plus maze | Yes | No | Yes |
|  | Open-field | Yes | Yes | Yes |
| – | Nest building | Yes | No | Yes |
|  | Marble burying | Yes | No | No |
|  | Startle Reactivity | No | No | No |
|  | Pre-pulse inhibition | No | No | No |
|  | Hot plate | No | No | No |

^1^Percentage is based on Stamberger et al., 2016 except for anxiety, which is based on Suri et al., 2017.

^2^For the phenotypes of mouse models, “Yes” indicates a statistical difference between mutant and control mice and “No” indicates that no statistical differences were detected between mutant and control mice.

^3^Dystonia only manifests as mild stiffness.

^4^*Stxbp1^tm1d/+^* and Vgat-cHet mice performed better than control mice at the age of 6–8 weeks.
